# LOCOPOTS: a LOw-COst high-throughput screening platform for in vitro POTatoes under abiotic Stress

**DOI:** 10.64898/2026.05.12.724622

**Authors:** Iñigo Saiz-Fernández, Lisney Alessandra Bastidas Parrado, Pavel Klimeš, Sanja Ćavar-Zeljković, José Ignacio Ruiz de Galarreta, Maria de la O Leyva-Pérez, Amaia Ortiz-Barredo, Lukáš Spíchal, Nuria De Diego

## Abstract

Potato crop is highly vulnerable to abiotic stresses like salinity and low nutrient availability. Rapid identification of stress-resilient genotypes is therefore essential for breeding, yet conventional phenotyping is often slow, space-demanding and expensive. We present LOCOPOTS — a LOw-COst high-throughput screening platform for *in vitro* POTatoes under abiotic Stress — which combines individual *in vitro* plant culture, low-cost RGB imaging and machine-learning-based automatic segmentation using a trained model of a convolutional neural network, based on U-Net architecture. LOCOPOTS enabled the automated extraction of growth, colour, and vegetation-index traits and demonstrated robust performance across independent phenotyping rounds. We screened 30 potato varieties under control, low-nutrient and saltinity conditions, identifying contrasting growth and physiological responses. Integrated traits such as final area and height, Area_AUC and height_AUC, together with GLI, Ch_ol_, cive and chlorophyll fluorescence parameters, discriminated genotype performance under stress. Metabolic profiling further revealed genotype-specific reprogramming in carbon and nitrogen metabolism under low nutrition and salt stress, including changes in fructose, myo-inositol, β-aminobutyric acid, γ-aminobutyric acid, proline, and certain polyamines, identifying them as specific chemical biomarkers of plant stress responses. LOCOPOTS provides a scalable, affordable and space-efficient platform for early screening of potato genetic diversity and identification of candidate traits associated with stress resilience.

## Introduction

Potato (*Solanum tuberosum* L.) is the fourth most cultivated crop in the world, and the leading non-cereal crop for human consumption (Johnson *et al*., 2025). Its high nutritional value, broad culinary use, and ability to produce large amounts of edible biomass per unit of water have made the potato a key component of global food security (Wijesinha-Bettoni and Mouillé, 2019). In addition, this crop possesses substantial intraspecific genetic variability, which has enabled its cultivation across contrasting agroclimatic regions and production systems (Slater *et al*., 2017). Potato is currently grown on more than 17 million ha worldwide (Food and Agriculture Organization of the United Nations (FAO)., 2024), spanning more than 100 countries (Rozentsvet *et al*., 2024), with an estimated global annual production of 300 million metric tonnes (Ojha *et al*., 2025). This data reflects both potato agronomic importance and its capacity to adapt to diverse environments (de Jesus Colwell *et al*., 2021; Rozentsvet *et al*., 2024).

Despite this broad adaptability, potato production remains highly vulnerable to abiotic stress. Although potatoes are among the most water-efficient crops, with one of the highest calorie-to-water ratios (Vreugdenhil *et al*., 2007), their shallow root systems make the crop highly sensitive to nutrient availability (Johnson *et al*., 2025). These characteristics have contributed to the intensive use of fertilisers in potato production systems, increasing production costs and environmental impacts (Li *et al*., 2019; Yang *et al*., 2022). Excessive fertilisation, together with irrigation practices and altered rainfall patterns, can contribute to soil salinisation, while climate change is increasing the frequency and intensity of drought, heat, flooding, and other extreme events (Chourasia *et al*., 2021). Consequently, it has been estimated that 20% of arable land and 33% of irrigated land are affected by salinity (Ojha *et al*., 2025), representing approximately 1128 Mha worldwide (Chourasia *et al*., 2021). Salinity and osmotic stress are particularly relevant for potato, which is considered a moderately salt-sensitive crop and can suffer severe reductions in tuber production of up to 60% under adverse water and salt conditions (Sanwal *et al*., 2022). Improving potato tolerance to abiotic stresses while enhancing nutrient use efficiency (NUE) is, therefore, a major objective for sustainable potato production.

Developing stress-resilient potato varieties, however, remains a slow and resource-intensive process. Conventional crop breeding is time-consuming, and identifying a new cultivar can take up to 10 years (Slater *et al*., 2017). A major bottleneck in breeding programs is the time required for plant phenotyping, as large numbers of genotypes must be evaluated across multiple environments, developmental stages, and growth conditions (Johnson *et al*., 2025). Even though the search and measurement of the most relevant traits can be performed using traditional classical methods, these methods are often destructive, labour-intensive, and limited in temporal resolution. High-throughput phenotyping (HTP) has emerged as a powerful alternative, enabling non-invasive, repeated, and quantitative assessment of plant growth and physiology at a larger scale (De Diego *et al*., 2017; Zdrazil *et al*., 2025).

HTP approaches rely on automated multi-sensor equipment to obtain accurate, non-destructive data from a large number of plants over a short period of time (Rozentsvet *et al*., 2024). In potato, image-based HTP has already been used to capture morphological and physiological responses to abiotic stress using RGB imaging, chlorophyll fluorescence, thermal imaging, and hyperspectral sensors (Abdelhakim *et al*., 2024). These authors developed a multi-sensor phenotyping protocol to monitor potato responses to single and combined stresses, although the study was conducted on potted plants and focused on a single cultivar. More recently, deep phenotyping, combined with multi-omics approaches, has revealed complex molecular and physiological signatures of potato responses to single and combined abiotic stresses by simultaneously phenotyping 150 potted plants of a single selected genotype across six growth conditions. These studies demonstrate the power of HTP for dissecting potato stress responses. However, they also illustrate a key limitation of current potato phenotyping platforms: their reliance on controlled-environment or greenhouse-based systems for potted plants, which require substantial cultivation space, expensive infrastructure and considerable handling time. This constraint limits the number of genotypes and treatments that can be screened, particularly in early breeding or pre-breeding stages. The limitation is especially relevant for potato because, compared with small model species such as *Arabidopsis*, its growth habit and canopy architecture rapidly increase the area required for large-scale experiments. Thus, there is a need for complementary phenotyping strategies that miniaturise experimental systems while retaining sufficient throughput and quantitative power to screen for potato genetic diversity across diverse growth conditions.

I*n vitro* culture offers an attractive solution to this bottleneck. Micropropagated potato plants can be grown under highly controlled conditions, require little space, and allow precise manipulation of medium composition, including nutrient availability and salt or osmotic stress, among other (Ugena *et al*., 2018; Nisler *et al*., 2024). *In vitro* culture has successfully been used to screen drought-tolerant potato varieties (Anithakumari *et al*., 2011), highlighting the potential of this technology. However, the integration of potato *in vitro* culture with automated image-based phenotyping remains limited.

Here, we present LOCOPOTS — a LOw-COst high-throughput screening platform for *in vitro* POTatoes under abiotic Stress. This system combines optimised *in vitro* culture in individual vessels with RGB-based image acquisition and automated trait extraction to quantify growth, morphology, and colour-derived indices in potato plantlets. Using a collection of 30 potato genotypes, we aimed to: (i) adapt *in vitro* culture conditions for reproducible individual plant growth and imaging; (ii) develop and validate a low-cost, RGB-based high-throughput phenotyping pipeline; and (iii) identify image-derived traits and biochemical markers associated with responses to nutrient limitation and salt stress. By reducing space requirements and infrastructure costs, LOCOPOTS provides a complementary phenotyping strategy for early-stage screening of abiotic stress resilience in potato breeding and pre-breeding programmes.

## Materials and methods

### Plant material and growth conditions

30 potato (*Solanum tuberosum* L.) varieties were used in this study, including commercial and non-commercial varieties originating from different geographical regions. These included Beltza from Spain, represented with black colour; BF-15 from France, represented in blue-green; Agria, Gala, Nicola, and Wega from Germany, represented in green; Ambition, Avano, Bafana, Bintje, Carlita, Desiree, El Mundo, Estima, Jaerla, Marfona, Melody, Monalisa, NU 633 Frisia, Primura, Rodeo, Sante, and Vivaldi from the Netherlands, represented in blue; Hermes from Austria, represented in yellow; Maris Piper from UK, represented in brown; Bermuda and Cara from Ireland, represented in violets; and Kennebec and Red Pontiac from USA, represented in orange and red, respectively (Supplementary Fig. S1).

The *in vitro* material was obtained from healthy, disease-free mother plants provided by NEIKER (Arkaute, Spain) and the Potato Research Institute (Havlíčkův Brod, Czech Republic). Plants were micropropagated following the method described by Zdrazil et al. (2025). Briefly, mono-nodal segments of approximately 7-10 mm, each containing one axillary bud, were excised from the mother plants, and leaves were removed. The nodes were then transferred to 7.7 × 7.7 × 9.7 cm magenta boxes (Sigma-Aldrich, Darmstadt, Germany) filled with 75 mL of full MS medium supplemented with 30 g L ¹ sucrose (8 g L ¹ agar, pH = 5.7). Between 9 and 12 nodes were transferred into each magenta box to maintain the genotype collection. Explants were kept in a Conviron CMP 6010 growth chamber (Conviron, Winnipeg, Canada) under controlled conditions with a constant temperature of 22 °C, light intensity of 80 µmol photons m^-2^ s^-1^, and a long-day photoperiod of 16 h light/ 8 h dark. Plants were grown until they had generated enough internodes (4 -5 weeks) to provide material for multiplication and subsequent non-invasive HTP screening method.

### In vitro conditions for HTP screening

Square plastic vessels (1.35 × 1.35 × 8.6 cm; Tintometer GmbH, Dortmund, Germany) were used for *in vitro* potato phenotyping. The air-tight caps were drilled four times each with a 2.5 mm bore to allow gas exchange. Both vessels and caps were sterilized by submersion in a 5% sodium hypochlorite solution overnight. The material was then rinsed twice with autoclaved water, and the vessels left into the flow box until drying. Immediately after, they were filled with 4.3 mL of growth medium. Control plants were grown on full-strength MS medium, whereas the stress treatments consisted of either nutrient limitation with 1/5 MS containing reduced sucrose (6 g/L) or salt stress with full-strength MS supplemented with 50 mM NaCl. Each plant was subdivided into a 7-10 mm mono-nodal section containing one leaf and an axillary bud. The two most basal and apical nodes were discarded. The leaves of the remaining nodes were excised, and one node was carefully inserted in the centre of each phenotyping vessel until approximately a third of the node remained submerged in the media. The vessels were then closed with the perforated caps, and the gap on the top of each cap was filled with autoclaved glass wool (VWR International s. r. o., Stříbrná Skalice, Czechia) to minimise contamination.

### RGB image acquisition

The vessels were kept in the growth chamber for 21 days (3 weeks) under defined growth conditions described above. *In vitro* plants were imaged daily using a custom-built phenotyping system designed to ensure standardised imaging conditions, as described by Zdrazil *et al*. (2025) (Supplementary Fig. S2). Briefly, the system provided a uniform background and fixed positioning of all components, using 3D-printed holders fabricated in-house. The imaging was performed using a Raspberry Pi Camera Module V2 and two strips of cool-white LED lights for homogenous illumination. For imaging, plant samples were arranged in seven vessels, which were placed into the custom 3D-printed holder designed to reduce lens distortion effects (e.g., fisheye distortion) (Supplementary Fig. S2. The holder was then positioned at a predefined location within the imaging system to ensure consistent alignment with the camera and background.

### RGB image processing and trait extraction

Three different approaches were used for RGB imaging analysis: manual analysis using ImageJ, semi-automatic analysis using MorphoAnalyser GUI v1.0.9.8. (PSI, Drásov, Czech Republic), and automatic analysis using an in-house developed ML-based Python pipeline. In the manual, plant height was calculated using ImageJ (https://imagej.nih.gov/ij/) by tracing a path along the stem from the lowest to the highest point. As a semiautomatic system, we determined morphological parameters, such as area, perimeter, height, width and compactness, using MorphoAnalyser GUI v1.0.9.8. (PSI, Drásov, Czech Republic). A “general plant mask” was generated from representative images by selecting and excluding specific colour classes. This mask was then applied to all images while maintaining acceptable segmentation accuracy. The plant mask settings were as follows: mean filter size = 3, colour formula = 0-((4*B)-(4*R)+(2*G)), threshold = -1.01176, min object size = 100 and minimum hole size = 50. In the automatic approach, RGB images were analysed using an in-house Python-based pipeline incorporating an ML model, as described below.

### Machine Learning Model for automatic RGB image analysis

#### Machine learning model and validation

A convolutional neural network based on the U-Net architecture was developed for plant image segmentation. The model was implemented in Python (https://www.python.org/) using the TensorFlow/Keras framework. The dataset consisted of 297 paired RGB images and corresponding binary masks, where the masks represented the plant area. All images were resized to a fixed resolution of 512 × 256 pixels with three colour channels (RGB).

The network architecture followed a standard encoder–decoder U-Net design. The encoder path consisted of successive convolutional blocks with 3 × 3 kernels, ReLU activation functions and He-normal initialisation, followed by max-pooling layers for downsampling. Dropout layers with rates ranging from 0.1 to 0.3 were included to reduce overfitting. The decoder path used transposed convolutions for upsampling, with skip connections linking corresponding encoder and decoder layers to preserve spatial information. The model was trained using the Adam optimiser and binary cross-entropy loss function, with accuracy as an additional evaluation metric. Training was performed for up to 20 epochs with a batch size of 16 and a 10% validation split. Early stopping based on validation loss was applied to prevent overfitting. The model achieved an accuracy above 99% on both the training/validation sets, indicating good performance. The trained model was saved and subsequently exported to the ONNX format for further deployment and interoperability. All codes are available at ZENODO (10.5281/zenodo.20122647), with a readme for better understanding.

#### Image preprocessing

Input RGB images were spatially cropped to isolate the region of interest containing plant-vessel regions. Each image was subsequently divided into seven vertical segments corresponding to individual vessels. To minimise edge artefacts, specific preprocessing steps were applied. For the first vessel, border pixels were manually masked to remove edge shadows. For the last vessel, vertical structural noise, such as dark lines along the vessel edge, was detected using the probabilistic Hough transform (Hough, 1962) and removed via inpainting. Each vessel image was resized to 256 × 512 pixels before further AI processing.

#### Deep Learning-Based Segmentation and Trait Extraction

Plants were segmented using the pretrained U-Net model. The model output was passed through a thresholding step with a threshold of 0.1 to produce a binary segmentation mask. Post-processing included removal of small artefacts, defined as contours with an area < 50 pixels, followed by morphological analysis.

For each segmented plant, the masked RGB pixels were extracted, and mean RGB values were computed. Colour composition was further analysed by assigning each pixel to the nearest predefined colour group using a KD-tree-based nearest-neighbour approach. The predefined colour groups represented biologically relevant colour clusters with the following R/G/B values: 115/110/50, 119/98/74, 124/140/65, 134/123/68, 145/157/72, 154/147/126, 160/143/110, 183/172/106, 30/45/14, 56/82/24, 85/108/24, and 98/86/85. These RGB values covered the colour spectrum observed across all potato genotypes and treatments. The number of pixels assigned to each colour group was recorded, generating a colour distribution profile for each plant. To enable comparison among plants of different sizes, pixel counts for each colour group were normalised to total plant area and expressed as relative abundance. These relative colour abundances (RA) were used for statistical analysis.

For structural analysis, binary masks were skeletonised using the Zhang–Suen thinning algorithm (Zhang and Suen, 1984). The resulting skeletons were analysed using the FilFinder2D library to extract the longest continuous path representing the main plant structure. To improve robustness against segmentation artefacts, discontinuities in the skeleton were detected, and gaps were bridged by drawing straight lines between the nearest endpoints. A semi-automatic validation step was included, allowing the user to visually inspect the original image, skeleton, and corrected skeleton, and to select one of the following options: (i) corrected length, (ii) original length, (iii) individual segment lengths, or (iv) manual input. Plant length was estimated as the number of non-zero pixels in the final skeleton representation.

For each vessel, the following parameters were recorded: plant identification, plant area and height (in Pixels), mean RGB values, pixel counts per colour group, and the relative abundance of each colour group with respect to the total plant area. The data evaluation was divided into two Python scripts: one dedicated to plant height estimation and the other to extracting the remaining phenotypic features. All results were exported into a structured CSV file. The mean RGB values per plant were then used to calculate additional colour-related traits, specifically 50 RGB-based vegetation indices, following the formulas shown in Supplementary Table S1 (Kior *et al*., 2024).

### Chlorophyll fluorescence imaging

At the end of the experiment, on day 21 after explant transfer to the phenotyping vessel, chlorophyll fluorescence (ChlF) imaging was performed to assess photosynthetic performance. For this purpose, the phenotyping vessels were fixed to standard plastic blue trays using double-sided adhesive tape and positioned in the PlantScreen^TM^ Compact system (PSI, Drásov, Czech Republic). The ChlF imaging system has an LED light panel and a high-speed charge-coupled device camera (720×560 pixels, 50 fps, 12-bit depth). Modulated light of a known wavelength was used to detect the fluorescence signal. Three types of sources were used: (1) PAM short-duration measuring flashes (33 μs) at 618 nm, (2) orange-red (618 nm) and cool-white (6,500 K) actinic lights with maximum irradiance of 440 μmol m^-2^ s^-1^, and (3) saturating cool-white light with maximum irradiance of 3,000 μmol m^-2^ s-1 (Sorrentino *et al*., 2021). The trays containing the phenotyping vessels were automatically loaded into the system’s light-isolated imaging cabinet, which had a top-mounted LED light panel. After the 30-minute dark-adaptation period, when the PSII reaction centres opened, the well plates were automatically transported to the ChlF imaging cabinet. A 5-second flash of light at 0.5 μmol m^-2^ s^-1^ intensity was applied to measure the minimum fluorescence (F_0_), followed by a saturation pulse of 0.8 s at 6,000 μmol m^-2^ s^-1^ to determine the maximum fluorescence (F_m_). Plants were relaxed in the dark for 3 s, then exposed to 70 s of cool-white actinic light to drive photosynthesis and measure the peak rise in fluorescence (F_p_). Additional saturation pulses were applied at 8, 18, 28, 48, and 68 s during actinic illumination for 3 min, corresponding to Lss1, Lss2, Lss3, and Lss4 states at 500 μmol m^-2^ s^-1^ constant photon irradiance to obtain the light-adapted initial fluorescence (F_0_’) and steady state-fluorescence yield (F_t_). A final saturating blue light pulse (6,000 μmol m^-2^ s^-1^, 0.8 s) was applied to measure the maximum fluorescence in the light-adapted state (F_m_’), and the level of fluorescence was measured just before the saturation pulse was considered the steady-state fluorescence in the light-adapted state (F_t_).

After measurement and parameter extraction, the PlantScreen™ Data Analyzer software (PSI, Drásov, Czech Republic) calculated several chlorophyll fluorescence-related parameters: maximum quantum efficiency of PSII (QY_max_ = F_v_/F_m_), steady-state PSII quantum yield (QY = F_m_’-F_t_/F_m_’), maximum efficiency of PSII in the light-adapted steady state (F_v_/F_m_ = F_m_’-F_0_’/F_m_’), steady-state non-photochemical quenching (NPQ = F_m_-F_m_’/F_m_’), coefficient of photochemical quenching in the steady-state estimate of the fraction of open PSII reaction centres PSII_open_/(PSII_open_+PSII_closed_) [qP = ((F_m_’- F_t_)/(F_m_’-F_0_’)], coefficient of non-photochemical quenching in steady state (qN= F_m_-F_m_’/F_m_), fraction of open PSII reaction centres [qL = (F_m_’-F_t_)/(F_m_’-F_0_’)×(F_0_’/F_t_)], and electron transport rate (ETR = QY×PAR×α×*f*). In the ETR calculation, *f* = 0.5 was used to account for equal partitioning of excitation energy between PSII and PSI, and α = 0.84 was used as the assumed leaf absorbance of photosynthetic tissues (Maxwell and Johnson, 2000).

### Targeted metabolomics

After phenotypic analysis of the 30 potato genotypes, selected varieties were used for targeted metabolomic analysis. Samples were collected 21 days after transfer into the phenotyping vessels. The bottom part of the original mono-nodal section was excised and discarded, and the remaining plant tissue was immediately frozen in liquid nitrogen, stored at –80 °C, lyophilised to complete dryness using a Martin Christ Beta 1-8 LDplus (Martin Christ, Germany), and homogenised in a mixer mill (Retsch, Germany). All homogenised samples were stored at –20°C until extraction.

Free polyamines and amino acids, as well as their acetylated forms, were analysed in lyophilised shoots using the method described by Ćavar Zeljković *et al*. (2024), with slight modifications. Briefly, 3–5 mg of plant material was extracted with 50% EtOH. Individual plants were used as independent biological replicates where possible; when biomass was insufficient, two plants were pooled. In all cases, the plant ID was recorded to enable accurate correlation with phenomics-derived traits.

From the extraction, three aliquots of 250 µL were prepared. The first two aliquots were used for the analysis of free amino acids and for the benzoyl chloride derivatisation of free polyamines, respectively. Free amino acids and acetylated metabolites were separated on a BEH AMIDE column, while derivatised free polyamines were separated on a reverse-phase BEH C18 column. Analytes were identified and quantified using a Nexera X2 UHPLC system (Shimadzu Handels GmbH, Kyoto, Japan) coupled with an MS-8050 mass spectrometer (Shimadzu Handels GmbH, Kyoto, Japan).

The third aliquot of 250 µL was used to quantify small organic acids and sugars using a slightly modified method of Lisec et al. (2006). Ribitol was added as an internal standard to each aliquot, and then evaporated to dryness under vacuum at 40°C. Samples were then methoximated with MeOX in pyridine (20 mg mL^-1^; 20 µL) for 1 h at 50 °C, followed by silylation with 30 µL MSTFA for 30 min at 40°C. The reaction mixture was transferred to a GC vial and analysed immediately. Trimethylsilyl derivatives were analysed by GC-MS using an Agilent Technologies 7890A gas chromatograph equipped with an HP-5MS UI capillary column (60 m × 0.25 mm × 0.25 µm) coupled to a 5975C mass selective detector. Helium was used as the carrier gas at 1.1 mL min^-1^, and samples were injected in splitless mode using an injection volume of 1 µL. The MS detector operated in EI mode at 70 eV with a scan range of 40–1000 amu. Compounds were identified using retention indices, authentic standards and library spectra (NIST/EPA/NIH), and quantified using external calibration curves.

### Statistical analysis

All statistical analyses were performed using the software RStudio version 2026.01.1+403. To validate the reproducibility of the developed phenotyping method, data from four independent experiments were visualised as violin plots, and statistical differences among experimental rounds were analysed using the non-parametric Kruskal-Wallis test.

Once the segmentation and trait extraction method had been established, phenotyping-related traits and metabolic data were analysed using principal component analysis (PCA) and correlation matrices to visualise data structure, assess relationships between phenotypic traits and metabolites, and identify the most informative parameters for subsequent analysis. PCA and correlation plots were generated using the R packages *factoextra* (Kassambara and Mundt, 2016, Preprint), *ggplot2* (Wickham, 2011), *grid* (Murrell, 2002), *ggrepel* (Slowikowski, 2016, Preprint), *ggnewscale* (Campitelli, 2019, Preprint), and *corrplot* (Wei and Simko, 2010, Preprint).

Selected traits and metabolites were then analysed using one-way or two-way ANOVA, depending on the experimental design. One-way ANOVA was used to compare genotypes under control conditions, whereas two-way ANOVA was used to test the effects of genotype, treatment, and their interaction. Statistical analyses and post hoc comparisons were performed using the *agricolae* package (de Mendiburu and Yaseen, 2006, Preprint).

## Results and discussion

Phenotyping of potato plants is often a time-consuming, space-demanding, and costly process (Slater *et al*., 2017; Rozentsvet *et al*., 2024; Johnson *et al*., 2025). To address these limitations, we developed a pipeline (Fig. 1) that reduces space through *in vitro* culture, lowers phenotyping costs by using affordable RGB sensors, and minimises image-processing time through automated segmentation and trait extraction. The study was divided into four main steps: i) *in vitro* potato phenotyping (Fig. 1A), ii) image evaluation (Fig. 1B), iii) phenotyping method validation (Fig.1C), and iv) biochemical analysis (Fig. 1D).

**Fig. 1.**
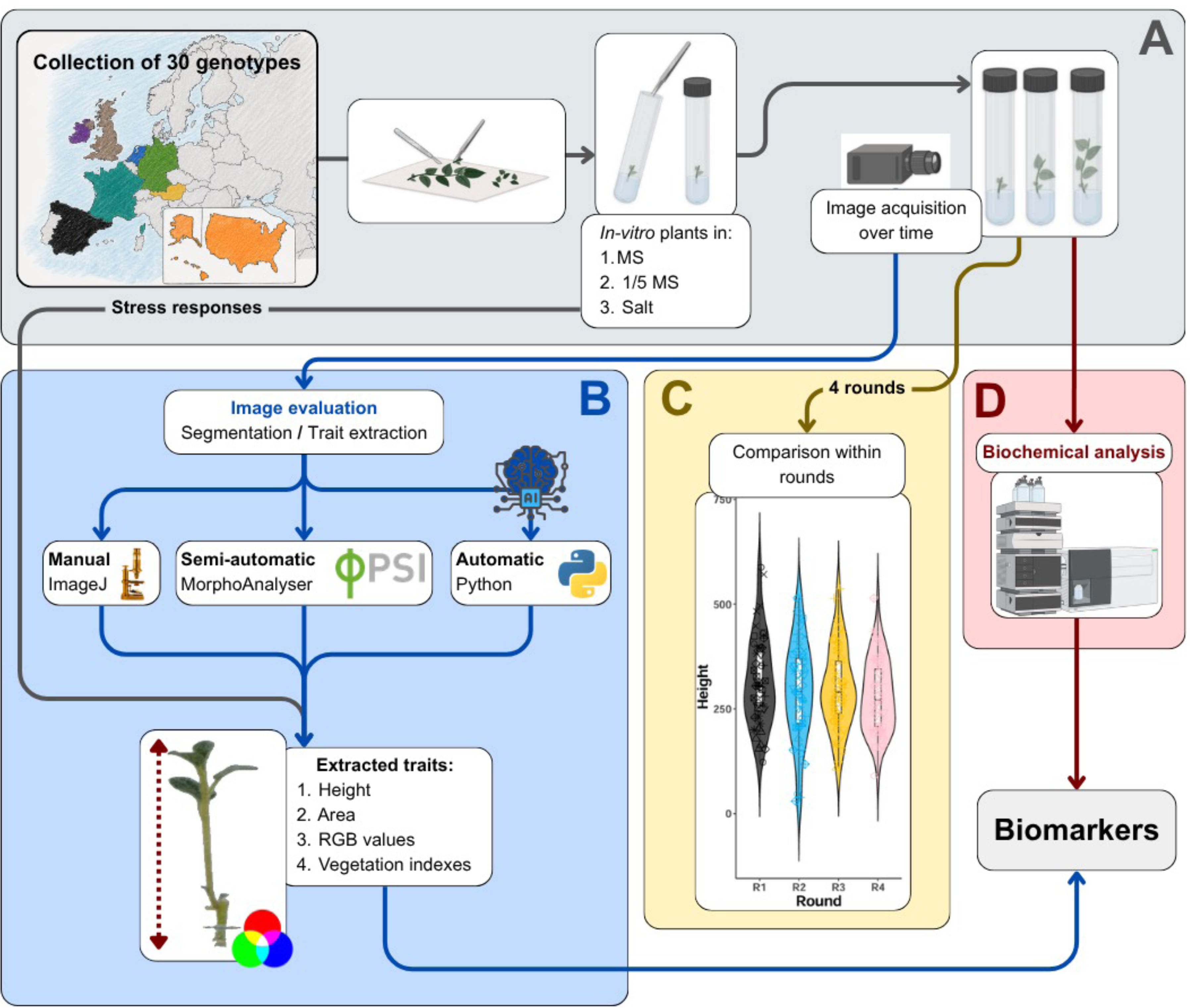
Workflow of the LOCOPOTS platform for *in vitro* phenotyping, image analysis and biochemical characterisation of potato genotypes under abiotic stress. (A) Thirty potato genotypes were propagated *in vitro* and grown under optimised growth conditions, including full-strength MS medium, low MS medium (1/5) and salt stress, followed by time-course image acquisition. **(B)** Images were processed using manual, semi-automatic and automatic approaches for segmentation and trait extraction, including ImageJ, MorphoAnalyzer and an in-house Python pipeline. Extracted traits included plant height, projected area, RGB values and vegetation indices. **(C)** Method reproducibility was assessed across four independent phenotyping rounds. **(D)** Biochemical analyses of selected metabolites were performed to identify stress-associated biomarkers.

### Optimisation of the in vitro *potato phenotyping setup*

The first challenge was to develop an *in vitro* setup suitable for image-based phenotyping of potato plants. Initially, we tested the cylindrical glass tubes routinely used for *in vitro* potato culture (Fig. 2A). However, the circular shape of the tubes caused substantial image distortion, making it unsuitable for accurate RGB imaging. Due to the limited options available for square-shaped glass tubes, we evaluated plastic alternatives (Fig. 2B). We selected square plastic vessels (1.35 × 1.35 × 8.6 cm) made of thicker plastic, with suitable dimensions and optical properties, originally designed for water quality testing (Tintometer GmbH, Dortmund, Germany). These vessels were not autoclavable and were therefore sterilised with sodium hypochlorite. This sterilisation treatment did not compromise vessel integrity or transparency, and adequate sterility was achieved.

**Fig. 2.**
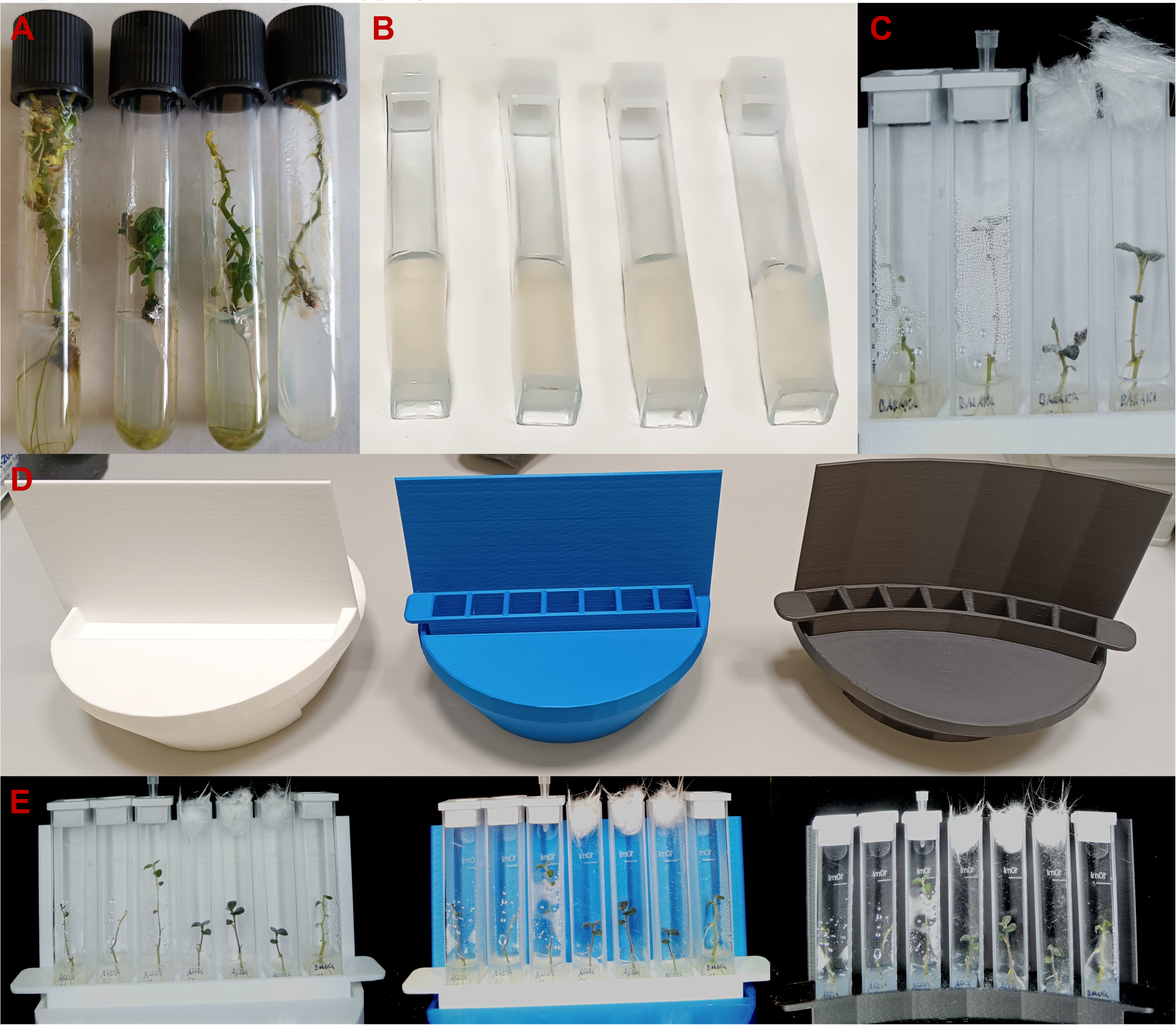
Optimisation of the *in vitro* potato growth setup for RGB-based phenotyping. **A)** Testing of RGB imaging using standard cylindrical tubes routinely used for in vitro potato culture. **B)** Testing of RGB imaging in square plastic vessels. **C)** *In vitro* growth of potato plants in square plastic vessels closed with airtight caps or covered with glass wool to allow gas exchange and reduce condensation. **D)** 3D-printed holders with different background colours — white, blue, and black— and angles to position the vessels for imaging, straight or with curvature, designed to facilitate RGB imaging and plant segmentation. **E)** Representative RGB images showing the effect of holder background colour and vessel-covering strategy on image quality. In each image, the right-hand vessel represents the perforated-cap setup, which reduced condensation and improved visibility for phenotyping.

During the first phenotyping test with potato plants grown in the square plastic vessels, it became evident that the standard airtight plastic caps did not allow any gas exchange. This affected plant morphology, resulting in elongated internodes and very small leaves (Fig. 2C). Moreover, limited gas exchange led to humidity build-up inside the vessels, causing strong condensation on the inner vessel walls and reducing imaging quality. Removing the caps and covering the vessels with glass wool restored normal plant morphology and reduced condensation (Fig. 2C). To limit the risk of contamination while maintaining gas exchange, the caps were perforated with four ventilation holes of 2.5 mm in diameter, which were then covered with sterilised glass wool. Plants grown using this setup showed morphology and size comparable to those grown with glass wool only, while maintaining acceptable condensation and a low risk of contamination.

Simultaneously, we tested 3D-printed holders with different background colours, including white, blue and black, to determine the most suitable configuration for plant segmentation (Fig. 2D-E). The holders not only presented different colours but also different geometries. Some holders were printed to align the vessels in a straight row, whereas others included a slight curvature to position the outer vessels closer to the camera and reduce edge-related optical distortion. The white holder provided the clearest plant recognition and facilitated segmentation (Fig. 2E). In addition, curved holders yielded the best segmentation images, as they reduced fisheye distortion at the border. Moreover, as shown in Fig. 2E, the perforated-cap setup was the best option for phenotyping *in vitro* potato plants, as demonstrated by the right-hand vessel in each RGB image, which showed reduced condensation and improved plant visibility.

### RGB image segmentation, trait extraction and validation

Once the setup for *in vitro* potato phenotyping was established, we evaluated image-processing alternatives, focusing on two main steps: plant segmentation and trait extraction. For plant segmentation, three approaches were used: i) ImageJ, a manual method in which each plant was analysed individually and only plant height was extracted; ii) MorphoAnalyzer, a semi-automatic method requiring manual definition of the plant mask and segmentation equation, which enabled the extraction of multiple traits, including plant height, area, compactness and width, and the pixels contributing of defined colour classes; and iii) in-house Python-based pipeline, an automatic method in which plant were identified and segmented without manual mask creation, allowing immediate extraction of traits such as plant height, area, relative and absolute growth rate (RGR and AGR) and the contribution to predefine colour classes.

To compare the reliability of the three different RGB image processing methods, ten randomly selected potato varieties grown under optimal conditions on full-strength MS medium were imaged across four independent rounds (Fig. 3). Final plant height and the accumulated height, calculated as the area under the growth curve (height_AUC) over the 21-day phenotyping period, were used to compare methods. The three methods were comparable and produced highly similar trends for both final plant height and height_AUC (Fig. 3D-I). All approaches identified Carlita as the slowest-growing variety within this subset, whereas Agria, Baraka, Estima, Gala, and Wega were among the fastest-growing genotypes. While final plant height provides a static measurement of plant development at a given stage, height_AUC integrates growth over the entire phenotyping period and, therefore, provides additional information on growth dynamics. The results suggest that both MorphoAnalyzer and the Python-based pipeline can reliably estimate plant height in *in vitro*-grown potato plants. However, the in-house pipeline, which uses a U-Net-based ML segmentation method, reduced post-processing time by eliminating manual mask creation and reducing the number of outliers. U-Net is a convolutional neural network architecture commonly used for biomedical and plant image segmentation, including several plant species (Bhagat *et al*., 2022; Deng and Wen, 2023; Zhu *et al*., 2026). Here, we trained a U-Net model to segment small potato explants grown *in vitro*, showing accurate segmentation and robust performance (99% accuracy) under our image conditions.

**Fig. 3.**
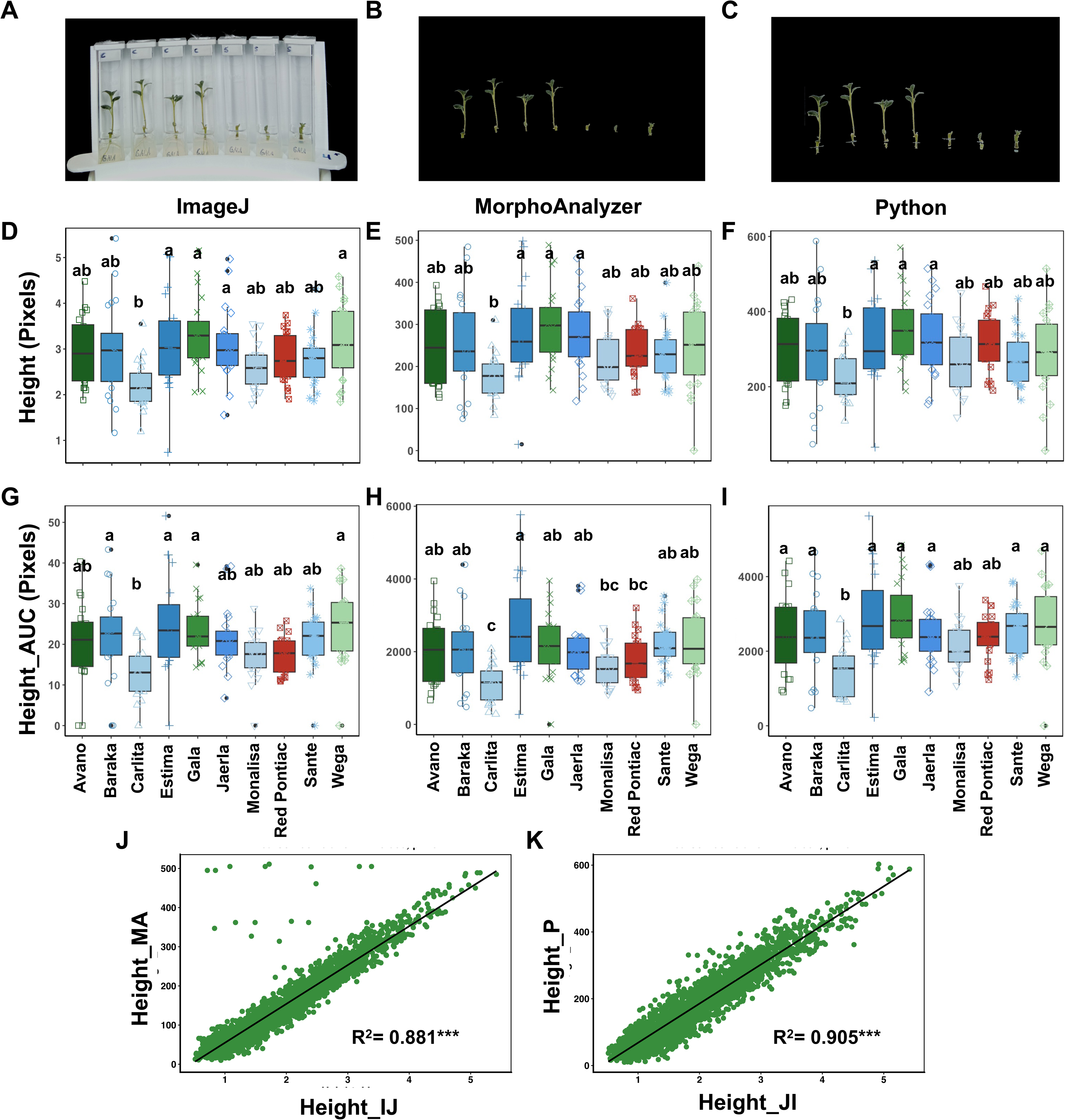
Optimisation and validation of plant segmentation for LOCOPOTS *in vitro* phenotyping method. Total Plant height quantified using three image-analysis approaches: ImageJ (IJ), as a manual reference method **(A)**; MorphoAnalyzer (MA), as a semi-automatic method **(B);** and an in-house Python pipeline based on a trained U-Net segmentation model (P), as an automatic method **(C).** Boxplots compare plant height **(D, E, and F)** or the accumulated height, calculated as the area under the growth Curve (AUC) **(G, H, and I),** at 21 days after explant transfer. Scatterplots show the correlation between ImageJ (IJ) and MorphoAnalyzer (MA)**(J)** or between ImageJ (IJ) and the Python/U-Net pipeline (P) **(K),** with Pearson’s correlation coefficient and significance indicated. Colours indicate genotype origin: green, Germany; blue, the Netherlands; and red, the USA. 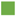 Germany; 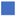 Netherlands; 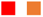 the USA.

A further comparison was performed by plotting manually calculated heights using ImageJ against those obtained with the semi-automated MorphoAnalyzer method and the automated Python-based pipeline (Fig. 3J -K). This analysis confirmed the robustness of both pipelines but also revealed several clear outliers in the MorphoAnalyzer dataset, visible in the upper-left region of the correlation plot (Fig. 3J). These points corresponded to images in which the top edges of the vessels were segmented as part of the plant, likely due to reflections, leading to a strong overestimation of plant height. Even though these data points could be removed during data cleaning, this would result in data loss, which may be particularly problematic when working with a limited number of replicates.

In a second step, we used the same dataset generated from four independent rounds, using the ten potato varieties shown in Fig. 3, to evaluate the reproducibility of the LOCOPOTS *in vitro* phenotyping methods (Fig. 4). The violin plots showed highly similar data distribution across the four rounds, with no significant differences between rounds according to the Kruskal-Wallis test(Fig. 4). Given that the rounds were performed several months apart, with more than a year between the first and last, the results demonstrate the robustness of the experimental setup and its suitability for phenotyping experiments conducted across multiple rounds over extended periods.

**Fig. 4.**
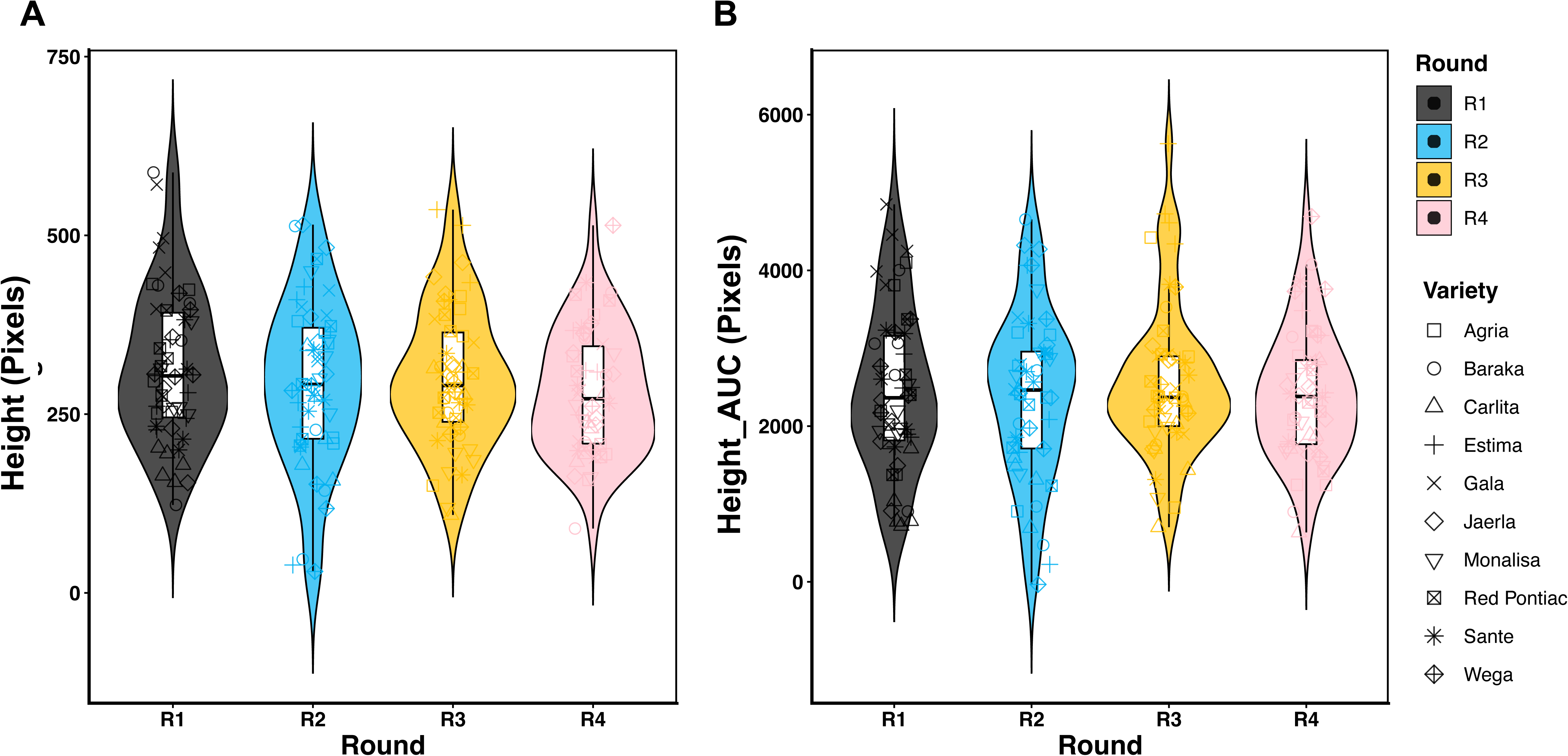
Reproducibility of the LOCOPOTS *in vitro* phenotyping method. Final plant height (A) and accumulated height, calculated as the area under the curve (AUC) (B) at 21 days after the explant transfer, from four independent rounds using ten potato genotypes grown under full MS medium and calculated using an in-house Python pipeline.

### LOCOPOTS for faster potato phenotyping

Once the LOCOPOTS pipeline had been developed, optimised and validated, we phenotyped 30 potato varieties from different origins and with different harvest classes: early, medium or late, as a proof of concept to assess whether genotype origin or harvest was associated with different growth and RGB-image-derived phenotypic traits. PCA was performed using all traits extracted from the RGB image of plants grown on full-strength MS medium for 21 days after explant transfer (Fig. 5A). The first two PCs accounted for 77.2% of the total variance (Fig. 5A). The biplot separated the varieties across the entire plot, but their distribution was not associated with their origin or harvest class (Fig. 5A). Moreover, growth-related traits, such as the final height, Height_AUC, the final area, and Area_AUC, clustered with defined colour classes and vegetation indices, and several of these traits were significantly correlated, as determined by Pearson correlation analysis (Supplementary Fig. S3A).

**Fig. 5.**
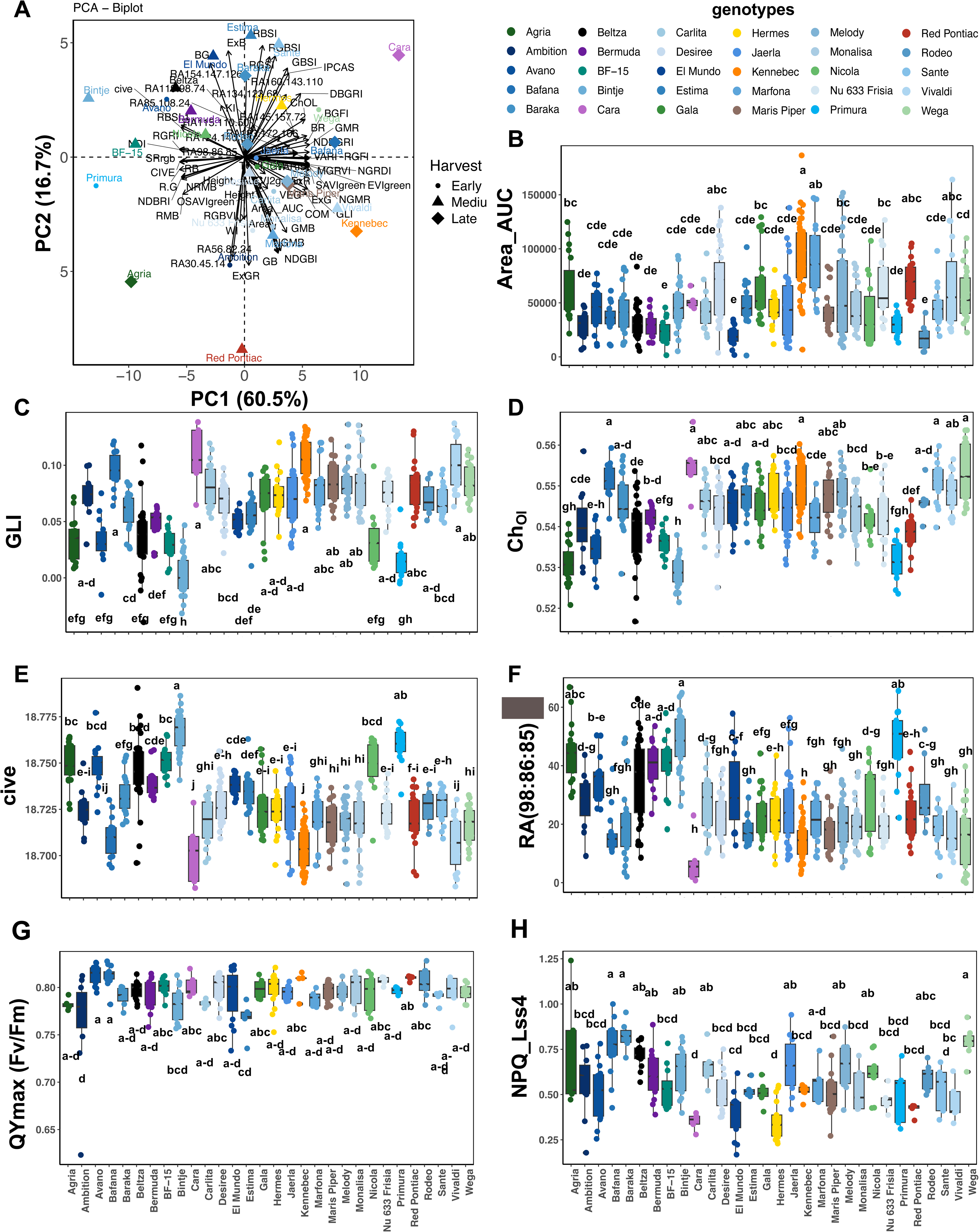
RGB Image-derived morphological and colour traits, and ChlF-derived traits of potato genotypes grown *in vitro* under control conditions. (A) Principal component analysis (PCA) performed using image-derived traits from 30 potato genotypes grown on full-strength MS medium for 21 days after explant transfer. The analysed features included four morphological traits — plant height, projected area, height AUC and area AUC — together with 15 segmentation-derived colour parameters and 50 colour-based vegetation indices. Representative genotype-specific distributions are shown for selected traits contributing to genotype separation, including Area_AUC **(B),** green leaf index (GLI) **(C),** chlorophyll index (Ch_ol_) **(D),** colour index of vegetation extraction (cive) **(E),** and the relative abundance (RA) of the RGB colour class 98:86:85 **(F),** the maximum efficiency of photosystem II (QY_max_) **(G),** and the non-photochemical quenching (NPQ_Lss4) (H). Different letters indicate significant differences according to Tukey’s HSD test following one-way ANOVA. Colours indicate genotype origin: 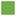 Germany; 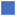 Netherlands; 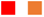 the USA; 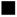 Spain; 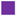 Ireland; 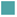 France; 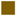 UK; and 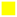 Austria.

Further statistical analysis was performed on representative traits that clustered together in the PCA and correlation matrix (Fig. 5A; Supplementary Fig. S3A). As an example of a growth-related parameter, we selected Area_AUC, calculated as the area under the growth curve (Fig. 5B). This parameter distinguished slow-growing varieties, such as BF-15, El Mundo, or Rodeo, from those with faster growth, such as Kennebec, Marfona, or Red Pontiac. Several vegetation indices derived from RGB images were positively correlated with plant growth parameters, such as the Green Leaf Index (GLI) or the chlorophyll index (Ch_ol_), whereas others, such as the colour index of vegetation extraction (cive), were negatively correlated (Supplementary Fig. S3A). GLI is an indicator of overall plant greenness and is often correlated with chlorophyll content and plant nutritional status, particularly nitrogen content (Hunt *et al*., 2013; Liu *et al*., 2023). Similarly, the Ch_ol_ index correlates with plant chlorophyll content (Ali *et al*., 2012; Riccardi *et al*., 2014). Both indices are useful to monitor plant health, as they tend to decrease in stressed plants (Ugena *et al*., 2018; Kior *et al*., 2024). In both cases, the highest values were observed in the well-growing varieties Kennebec and Marfona, as well as in the genotype Cara (Fig. 5C-D). However, Cara was in a different quadrant of the biplot due to its average growth (Fig. 5A and 5 B), indicating complementary information provided by these traits. In contrast, Bintje showed the lowest values of GLI and Ch_ol_, an average biomass accumulation and the highest values of cive. This index negatively correlated with plant height and area and was reduced in varieties with the highest GLI and Ch_ol_, such as Kennebec or Cara (Fig. 5E, Supplementary Fig. S3A).

Cive is usually employed to separate plants from the background in top-view or aerial RGB images, as it enhances soil, ground, and non-photosynthetically active areas of the image (Beniaich *et al*., 2019; Benaissa *et al*., 2024; Kior *et al*., 2024). Since the RGB images for the calculation of this index have already been segmented before its calculation, cive is most likely capturing the brown-grey colour associated with senescence or necrotic tissues, or the accumulation of other nongreen-related pigments, as its value decreases with increasing green colour classes, such as the relative abundance (RA) of the medium-dark green colour 124:140:60, or the indices GLI and Ch_ol_. Moreover, cive was positively correlated with the colour class 98:86:85, which represents a dark green/grey region of the RGB spectrum. Bintje and Primura showed the highest relative abundance of this colour class, whereas Kennebec and Cara showed the lowest (Fig. 5F). This parameter highlighted darker-coloured varieties, which may indicate differences in pigment composition or the accumulation of nitrogen-rich compounds (Farhan *et al*., 2024). The accumulation of certain nitrogen-related compounds under excess nitrogen could limit plant growth, beyond nutrient availability, causing these compounds to accumulate in source tissues rather than being used to support newly growing tissues (Saiz-Fernández *et al*., 2017). Altogether, these results indicate that LOCOPOTS can efficiently extract a broad set of RGB-derived morphological and colour traits sufficient to phenotype potato genotypes based on their *in vitro* growth conditions. In addition, colour classes and vegetation indices provide indirect information on plant health status and potential biochemical variation, supporting their use as complementary traits for rapid genotype characterisation.

The data from the RGB sensor were complemented by non-destructive ChlF imaging to evaluate the photosynthetic performance of the potato varieties under control conditions (Fig. 5G and H, Supplementary Fig. S3). The first two PCs accounted for 65.5% of the total variance and primarily separated potato varieties with higher photosynthetic efficiency from those with lower efficiency (Supplementary Fig. 3B). Moreover, the correlation matrix showed that fluorescence-related parameters associated with better photosynthetic performance were positively correlated with each other and with RGB-derived traits, such as height and area, whereas they were negatively correlated with non-photochemical quenching parameters (Supplementary Fig. S3C). For this reason, we selected one representative parameter from each cluster for further comparison among genotypes: QY_max_ (F_v_/F_m_), as an indicator of maximum PSII efficiency, and NPQ_Lss4, as an indicator of energy dissipation (Fig. 5G-H). QY_max_, therefore, reflects the functional status of the photosynthetic apparatus (Zhao *et al*., 2021). Higher values of this parameter are typically found in plants with greater carbon assimilation capacity and greater potential for vegetative growth (Wang *et al*., 2019). Accordingly, fast-growing varieties such as Kennebec or Red Pontiac showed consistently high QY_max_ values (Fig. 5G). However, other genotypes, such as Avano and Bafana, had the highest QY_max_ values despite not showing the highest Area_AUC values. This suggests that high maximum PSII efficiency alone does not fully explain growth performance and that other physiological processes may limit biomass accumulation in these genotypes. Interestingly, Avano and Bafana also showed high NPQ_Lss4 values, together with Wega (Fig. 5H). NPQ estimates the fraction of absorbed light energy that is dissipated as heat rather than used for photochemistry (Bassi and Dall’Osto, 2021). The coexistence of high QY_max_ and high NPQ_Lss4 in some genotypes suggests that their PSII reaction centres retained high maximum efficiency, but that, under the light conditions used in this study, a larger fraction of absorbed light energy was dissipated as heat rather than used for photochemistry or downstream carbon assimilation (Maxwell and Johnson, 2000). Importantly, this increased energy dissipation occurred without apparent impairment of the maximum capacity of PSII, as indicated by their high QY_max_ values. Consistent with this interpretation, Lehretz et al. (2022) showed in potato that altered NPQ dynamics did not affect PSII efficiency-related parameters under low or normal light conditions but could limit CO_2_ fixation and reduce growth under specific light regimes. This could partially explain why these genotypes did not show the highest growth performance, since sustained or slowly relaxing NPQ can limit CO assimilation and biomass accumulation.

### LOCOPOTS enables rapid screening of potato responses to low nutrient availability

Next, we tested the ability of LOCOPOTS pipeline to identify potato varieties with contrasting responses to defined abiotic stress conditions. As a first stress-related application, we analysed growth and phenotypic changes under low-nutrient availability using 1/5 MS medium with reduced sucrose content. We first evaluated changes in RGB imaging-derived traits, colour classes and vegetation indices across the 30 genotypes (Fig. 6). To facilitate visualisation and interpretation, traits were expressed relative to their corresponding values under optimal growth conditions and were used for trait selection and genotype classification under low-nutrient conditions. In the PCA biplot, the first two PCs accounted for 58.1% of the total variance and distributed the genotypes across the four quadrants (Fig. 6A). PC1 mainly separated Bintje from the other varieties due to higher values of specific colour indices, whereas PC2 separated the varieties according to their growth responses under low-nutrient conditions (Fig. 6A). For example, under 1/5 MS, Bintje showed the tallest plants and largest projected area, whereas varieties such as Nicola showed reduced biomass accumulation (Fig. 6B). The height and area-related traits were further analysed using two-way ANOVA. Although a significant genotype × treatment interaction was detected, post hoc analysis did not identify significant differences between most genotypes and their corresponding controls grown under optimal conditions. However, Bintje increased final plant height by 20% and Area_AUC by 31% (Fig. 6B-C). Conversely, BF-15 and Nicola showed reductions of 54% and 48% in plant height, and 63% and 66% in Area_AUC, respectively. These results indicated that Area_AUC was a more sensitive parameter for capturing dynamic growth responses in plants under low nutrition.

**Fig. 6.**
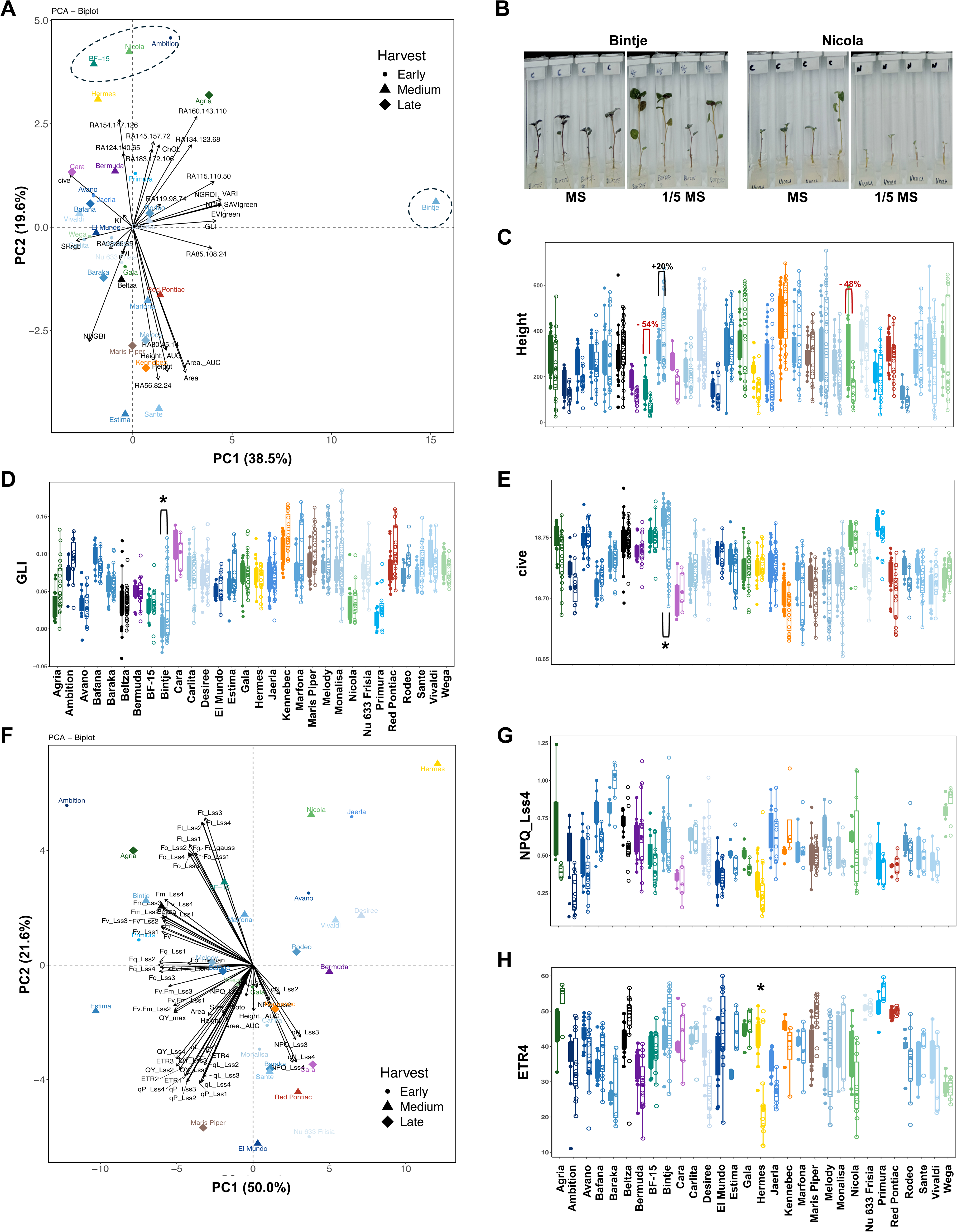
RGB-derived morphological and colour traits, and ChlF-derived fluorescence parameters, of potato varieties grown *in vitro* under low-nutrient conditions. (A) Principal component analysis (PCA) performed on morphological and colour-related traits extracted from RGB images and expressed as the ratio between plants grown on 1/5 MS medium and those grown on full-strength MS medium, 21 days after explant transfer. **(B)** Representative images of contrasting genotypes under low-nutrient conditions. Representative genotype-specific distributions are shown for selected RGB-derived traits contributing to genotype separation, including final plant height **(C)**, green leaf index (GLI) **(D)**, and colour index of vegetation extraction (cive) **(E)**. Asterisks indicate significant differences relative to the corresponding control plants grown on full-strength MS medium, according to Tukey’s HSD test following two-way ANOVA with genotype and treatment as factors. Filled boxplots and symbols represent plants grown on full-strength MS medium, whereas open boxplots and symbols represent plants grown on low-nutrient medium. **(F)** PCA performed on fluorescence-related traits extracted from ChlF imaging. Representative genotype-specific distributions are shown for selected fluorescence parameters: NPQ_Lss4 **(G)** and ETR4 **(H)**. Colours indicate genotype origin: 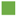 Germany; 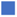 Netherlands; 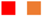 the USA; 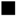 Spain; 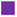 Ireland; 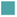 France; 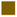 UK; and 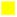 Austria.

Similar patterns were observed for the vegetation indices GLI and cive, which were significantly correlated with biomass-related parameters in plants under low-nutrient conditions, with positive and negative correlations, respectively (Supplementary Fig. S4A). Interestingly, Bintje was the only genotype showing significant differences in both GLI and cive under low nutrient conditions compared with its control plants grown under optimal conditions (Fig. 6D-E). This is consistent with previous studies highlighting GLI for identifying plant tolerance or susceptibility to stress (Ugena *et al*., 2018), as it reflects the capacity of plants to maintain leaf greenness under stress conditions, which is often associated with chlorophyll content and nutritional status (Hunt *et al*., 2013; Liu *et al*., 2023). Moreover, our results also suggest that cive may be an informative colour-derived marker for detecting stress-related changes after plants segmentation, particularly in systems where the background does not resemble soil or substrates.

ChlF imaging-extracted parameters were also analysed by PCA, revealing a genotype distribution that differed from that obtained using RGB-related traits (Fig. 6F). Potato varieties were mainly separated into three groups. One group, located in the upper-left quadrant, included two subgroups: Bintje and Primura, which showed higher values of fluorescence parameters such as F_v_ and F_m_, and Agria and BF-15 with higher values of F_0_ and Ft. All four varieties also tended to reduce the non-photochemical quenching-related parameters such as NPQ_Lss4 (Fig. 6F-G). Increased F_0_ can reflect reduced efficiency of excitation energy transfer from the light-harvesting antenna to PSII reaction centres or partial impairment of PSII reaction centres (Rosenqvist and van Kooten, 2003). Moreover, nitrogen-dependent changes in fluorescence parameters have previously been reported in potato, with F_m_, F_v_, and F_v_/F_m_ responding to nitrogen supply and genotype, and F_v_ and F_m_ showing a positive association with chlorophyll content (Mauromicale *et al*., 2006). Interestingly, BF-15 tended to have the highest F_0_ values under low nutrient availability, along with a marked reduction in biomass and GLI, suggesting that this genotype may require greater nutrient availability to maintain growth and photosynthetic performance. The combination of high F0, reduced biomass, and lower GLI may indicate sensitivity to low nutrition, although no changes in QY_max_ were observed for BF-15.

A second group located in the upper-right quadrant included Hermes, Nicola and Jaerla, which appeared to be among the varieties most affected by low nutrient availability (Fig. 6A). Specifically, Hermes and Nicola were among the genotypes showing the strongest reductions in growth-related traits (Fig. 6C; Supplementary Fig. SX). These three varieties also had reduced QY_max_ and lower values of F_v_/F_m_, QY, qP, qL, and ETR along the light-response curve (Fig. 6F-H), parameters that were positively correlated with plant growth (Supplementary Fig. S4B). This suggests that Hermes, Nicola, and Jaerla may be more sensitive to low nutrient availability at the level of PSII photochemistry and photosynthetic electron transport (Rosenqvist and van Kooten, 2003). Moreover, these results support the usefulness of combining fluorescence traits with RGB-derived traits to detect genotypes with contrasting responses to low nutrient availability.

The third group included varieties such as Baraka, Cara, Sante, Red Pontiac, Kennebec or Gala, which showed higher values of non-photochemical quenching-related parameters, according to the PCA (Fig. 6F). This response may indicate greater energy dissipation under low nutrient availability, suggesting that these genotypes activate photoprotective mechanisms to avoid excess excitation pressure when nutrient limitation restricts growth or downstream carbon metabolism. However, it is worth noting that ChlF parameters in stressed plants did not differ significantly from those under optimal conditions for most varieties. Only Hermes showed a significant reduction in ETR4 under low-nutrient conditions, suggesting that this genotype may be particularly sensitive to nutrient limitation at the level of photosynthetic electron transport.

### Low nutrient responses are associated with genotype-specific changes in carbon- and nitrogen-related metabolites

To better understand further the contrasting morphology and physiology observed in *in vitro* potato plants, we selected four genotypes with different responses under low nutrient availability. The selection included BF-15, which showed one of the strongest reductions in biomass under low nutrient availability and high F_0_ and F_t_ values; Nicola, which showed reduced biomass and increased non-photochemical quenching; Gala, which showed almost no visible changes in biomass and maintained fluorescence performance; and Bintje, which showed the highest biomass accumulation under low nutrient availability and higher F_m_ and F_v_ values (Fig. 7). These four varieties also displayed contrasting metabolic profiles.

**Figure 7.**
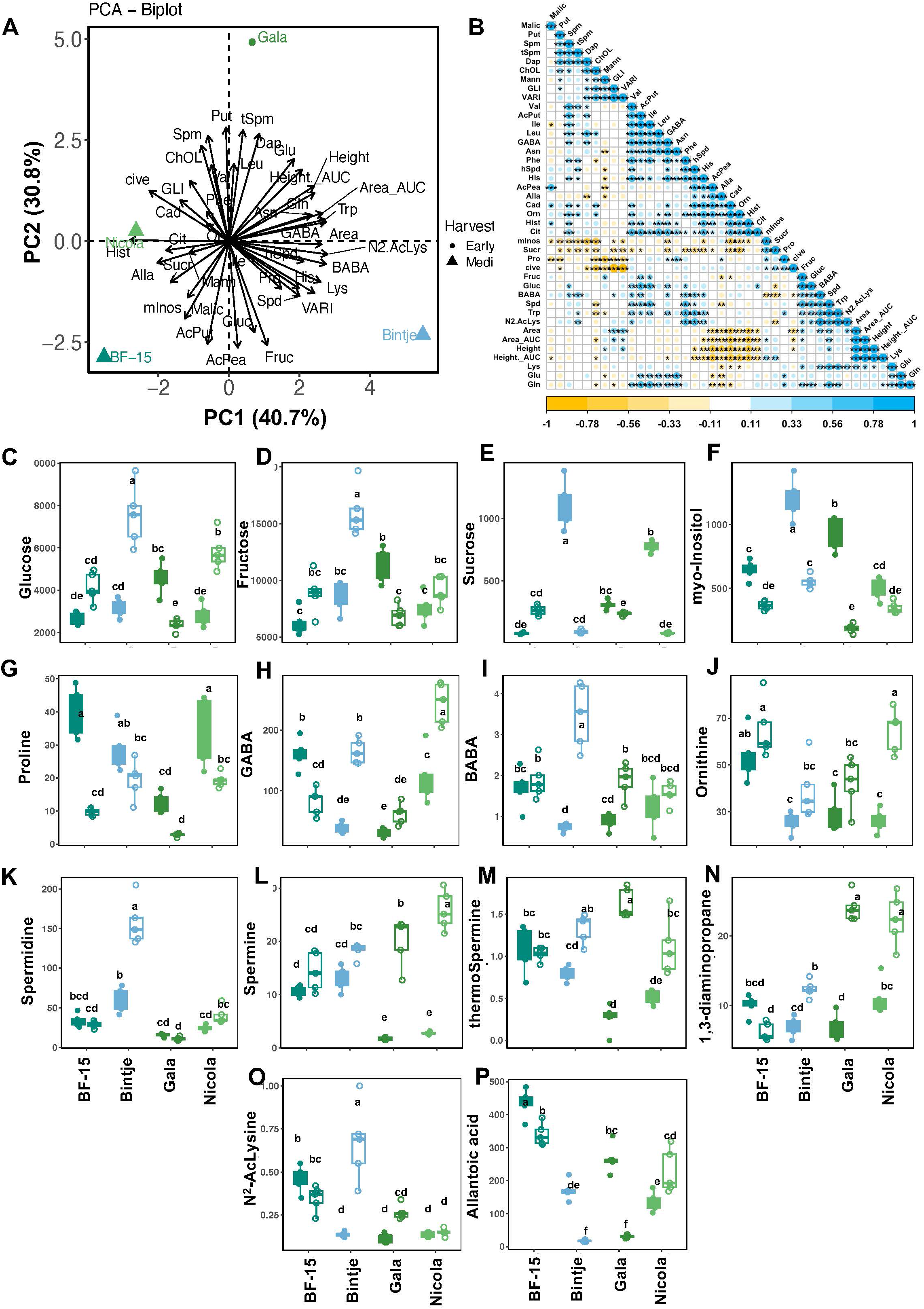
Biochemical changes in selected potato varieties grown *in vitro* under low-nutrient conditions. **(A)** Principal component analysis (PCA) performed using sugars, free amino acids, free and acetylated polyamines, and other stress-related metabolites, together with representative RGB-derived traits. Values were expressed as the ratio between plants grown on 1/5 MS medium and those grown on full-strength MS medium, 21 days after explant transfer. **(B)** Correlation matrix showing relationships between metabolite content and selected RGB-derived traits. Representative metabolites contributing to genotype separation are shown, including glucose (Gluc) **(C)**, fructose (Fruc) **(D)**, sucrose (Suc) **(E)**, myo-inositol (mInos) **(F)**, proline (Pro) **(G)**, γ-aminobutyric acid (GABA) **(H)**, β-aminobutyric acid (BABA) **(I)**, ornithine (Orn) **(J)**, spermidine (Spd) **(K)**, spermine (Spm)**(L)**, thermospermine (tSpm) **(M)**, 1,3-diaminopropane (Dap) **(N)**, N2-acetyllysine (N2AcLys) **(O)**, and allantoic acid (Alla) **(P)**. Different letters indicate significant differences according to Tukey’s HSD test following two-way ANOVA with genotype and treatment as factors. Filled boxplots and symbols represent plants grown on full-strength MS medium, whereas open boxplots and symbols represent plants grown on low-nutrient medium. Colours indicate genotype origin: 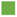 Germany; 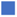 Netherlands, 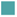 France. Additional metabolite abbreviations: acetylphenylethylamine (AcPea), acetylputrescine (AcPut), asparagine (Asn), cadaverine (Cad), citrulline (Cit), glutamic acid (Glu), glutamine (Gln), histamine (Hist), histidine (His), isoleucine (Ile), homospermidine (hSpd), leucine (Leu), lysine (Lys), malic acid (Malic), mannitol (Mann), phenylalanine (Phe), putrescine (Put), tryptophan (Trp), and valine (Val).

PCA, in which PC1 and PC2 explained 71.5% of total variance, indicated that, under low nutrient availability, enhanced growth was associated with higher accumulation of monosaccharides, particularly glucose and fructose, together with reduced sucrose levels, as observed in Bintje, especially regarding fructose (Fig. 7A, C-E). The accumulation of carbohydrates, particularly reducing sugars such as fructose and sucrose, together with other molecules, including the polyols mannitol and myo-inositol, or the amino acid proline (Pro), can contribute to osmotic adjustment under nutrient stress (Hermans *et al*., 2006; Singh *et al*., 2026, Preprint). For instance, potassium deficiency has been described to disrupt cellular ionic and osmotic balance (Zhu *et al*., 2020), and soluble sugars may help compensate for the osmotic role of missing ions while also contributing to charge balance and carbon redistribution (Cakmak *et al*., 1994; Hermans *et al*., 2006). Among the sugars measured here, fructose appeared to be the most informative marker under our conditions, due to its high accumulation in Bintje and its positive correlation with plant area (Fig.7B).

Under low nutrient availability, we expected Pro and myo-inositol to accumulate as compatible solutes. However, both metabolites decreased significantly in specific varieties: myo-inositol was reduced in BF-15, Bintje, and Gala, whereas proline was reduced in BF-15 and Nicola (Fig.7F-G). Interestingly, Bintje and Gala showed the highest myo-inositol levels under full-strength MS medium. Myo-inositol is synthesised from glucose-6-phosphate through the myo-inositol phosphate synthase (MIPS E.C. 5.5.1.4) pathway (Geiger and Jin, 2006; Roychowdhury *et al*., 2025). Therefore, the higher sucrose availability in full-strength MS may favour carbon flux towards inositol metabolism. In Bintje and Gala, elevated myo-inositol under optimal growth conditions may reflect excess carbon availability or altered osmotic/carbon status rather than improved growth performance. This interpretation is particularly relevant for *in vitro* culture, where sucrose provides both metabolic energy and contributes to the osmotic potential of the medium (Yaseen *et al*., 2013). The response of Bintje was especially notable: this variety reduced myo-inositol levels under 1/5 MS medium while improving growth. Although myo-inositol is generally associated with osmoprotection and stress signalling (Valluru and Van den Ende, 2011), its genotype-dependent accumulation may indicate different carbon-use strategies under low nutrient availability. In addition, myo-inositol catabolism has been linked to low energy/nutrient responses through the regulation of myo-inositol oxygenase genes, providing a possible mechanism by which free myo-inositol pools may decrease under nutrient limitation (Alford, 2012). Moreover, exogenous sucrose application has been shown to reduce the expression of certain myo-inositol oxygenase genes (Alford, 2012), further supporting a connection between sucrose availability and inositol metabolism. Together, these results suggest that sucrose availability can shape carbon-use strategies and influence the *in vitro* growth performance of specific potato genotypes. This highlights the importance of further optimising the composition of culture media for genotype maintenance and phenotyping, particularly in potato genebanks.

Low nutrient availability also modified the content of γ-aminobutyric acid (GABA) and β-aminobutyric acid (BABA). GABA has been reported to modulate nutrient uptake, carbon/nitrogen balance, and stress-related gene expression (Podlešáková *et al*., 2019; Jalil *et al*., 2025). Regarding BABA, its metabolism is not fully understood, although it has been detected endogenously in several plant species (Thevenet *et al*., 2017; Ćavar Zeljković *et al*., 2024). The Bintje-specific increase in BABA may reflect activation of a stress-priming mechanism under low nutrient availability. BABA is a naturally occurring non-protein amino acid whose levels can increase under stress and whose perception is linked to broad-spectrum priming through hormone-associated pathways, including abscisic acid- and ethylene-related responses (Thevenet *et al*., 2017; Virág *et al*., 2024). Therefore, BABA accumulation in Bintje may help maintain growth under low-nutrient conditions by modulating metabolism and stress responses (Virág *et al*., 2024). Together with fructose accumulation, BABA may represent an additional candidate marker associated with better *in vitro* performance under low nutrient availability.

The arginine and ornithine (Orn) pathways were also altered under low nutrient availability and influenced polyamine metabolism (Fig. 7). Orn levels were higher in varieties that showed the greatest reduction in growth under low nutrient availability. However, spermidine (Spd) accumulated significantly only in Bintje, whereas other genotypes accumulated spermine (Spm), thermospermine (tSpm), or 1,3-diaminopropane (Dap) (Fig. 7K-N). Spd accumulation in plants under stress conditions has been associated with better growth and stress tolerance (Marchetti et al, 2018). Moreover, exogenous Spd application has been linked to improved photosynthesis and more selective nutrient uptake under stress conditions, including nutrient imbalance associated with salinity (Xue *et al*., 2025; Rashid *et al*., 2026). Thus, the specific accumulation of Spd in Bintje may contribute to its improved growth under low nutrient availability, potentially by supporting photosynthetic performance, stress protection, or nitrogen remobilisation.

Finally, Bintje was the only variety showing increased *N^2^*-acetyl-lysine (N2-AcLys) content, whereas both Bintje and Gala had the lowest allantoic acid (Alla) levels. N2-AcLys is an acetylated amino acid derivative that may reflect changes in amino acid turnover and lysine metabolism. In Arabidopsis, N2-AcLys accumulation has been related to stress conditions (Ćavar Zeljković *et al*., 2024). Its specific accumulation in Bintje suggests that low nutrient availability may trigger genotype-specific adjustments in nitrogen-containing metabolites to better perform under these conditions. However, Alla is an intermediate of purine catabolism and is linked to nitrogen recycling and remobilisation in plants. The strong reduction of Alla in Bintje and Gala may therefore indicate altered purine-derived nitrogen metabolism, possibly reflecting a reduced reliance on ureide-associated nitrogen remobilisation or a shift in nitrogen allocation under low nutrient availability (Casartelli *et al*., 2019). Together, these results indicate that low nutrient availability triggered genotype-dependent reprogramming of carbon- and nitrogen-related metabolism in potato, with Bintje showing a distinctive metabolic profile characterised by the accumulation of fructose, BABA, Spd, and N2-AcLys, along with reduced levels of myo-inositol and Alla.

### LOCOPOTS enables rapid screening of genotype-dependent responses to salt stress

Lastly, we tested the ability of LOCOPOTS to detect genotype-dependent responses to salinity (Fig. 8), one of the most damaging environmental stresses on potato production (Sanwal *et al*., 2022). Salt stress was imposed using 50 mM NaCl, a relatively mild treatment intended to detect early or moderate differences in *in vitro* growth performance rather than severe stress injury. PCA of RGB imaging-derived traits from the 30 varieties under salt stress showed that PC1 and PC2 explained 52.1% of the total variance and separated them into four main groups (Fig. 8A). The first group, located in the upper-right quadrant, included varieties such as El Mundo, Baraka, Bermuda, or Rodeo, which maintained growth levels similar to those observed under non-stress conditions (Fig. 8A, C-D). Interestingly, most varieties in this group had medium or small sizes under control conditions (Fig. 8B-D). The slower growth may help the plants to regulate the stress, and when salt is applied, they are able to induce an adaptive response with a more “conservative” growth response during the first salt stress phase (osmotic stress) for better long-term survival because they avoid the rapid accumulation of toxic ions in their tissues (Munns and Tester, 2008). This suggests that resistance to environmental stresses is often driven by a conservative growth strategy in which plants may prioritise secondary metabolism at the expense of primary metabolism and plant development, preparing them for stress conditions (Pandit *et al*., 2024). A second group was represented by Bintje, which performed well under salinity. This genotype showed no major changes in Area_AUC and a moderate reduction in the final plant height (Fig. 8A-D). A third group, located in the bottom-right quadrant of the biplot, included genotypes such as Bafana, Jaerla, Gala, Monalisa, or Wega, which reduced growth under salinity. The fourth group, including BF-15 and Nicola, also showed reduced growth and was associated with a relative accumulation of the yellowish colour class 124:147:65 (Fig. 8A-B, Supplementary Fig. 5A). Among the morphological and colour-related traits, final plant height showed the clearest genotype-dependent differences, suggesting that it may be one of the most informative morphological markers for *in vitro* potato screening under moderate salinity.

**Fig. 8.**
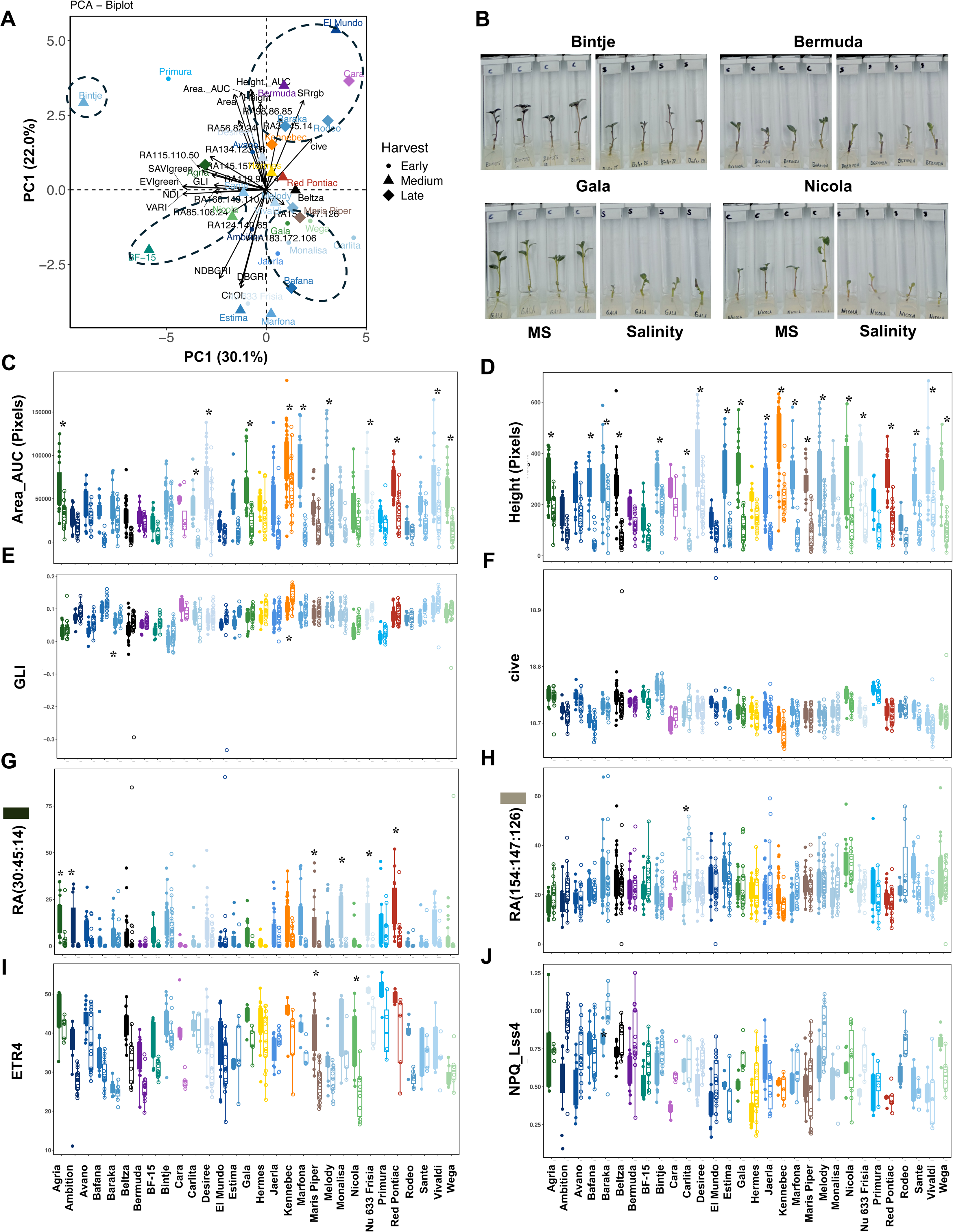
RGB image-derived morphological and colour-derived traits, and fluorescence parameters of potato varieties grown *in vitro* under salinity. (A) Principal component analysis (PCA) performed on morphological and colour-related traits extracted from RGB images and expressed as the ratio between plants grown on full-strength MS medium supplemented with 50 mM NaCl and those grown on full-strength MS medium, 21 days after explant transfer. **(B)** Representative images of contrasting genotypes under salt stress conditions. Representative genotype-specific distributions are shown for selected RGB-derived traits contributing to genotype separation, including accumulated projected area, calculated as the area under the curve (Area_AUC) **(C)**, final plant height **(D)**, green leaf index (GLI) **(E)**, colour index of vegetation extraction (CIVE) **(F)**, and the relative abundance of the dark-green RGB colour class 30:45:14 **(G)** and the earth-toned/khaki RGB colour class 154:147:126 **(H)**. Asterisks indicate significant differences relative to the corresponding control plants grown on full-strength MS medium, according to Tukey’s HSD test following two-way ANOVA with genotype and treatment as factors. Filled boxplots and symbols represent plants grown on full-strength MS medium, whereas open boxplots and symbols represent plants grown under salt stress. Representative genotype-specific distributions are shown for selected fluorescence parameters: ETR4 **(I)** and NPQ_Lss4 **(J)**. Colours indicate genotype origin: 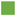 Germany; 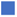 Netherlands; 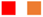 the USA; 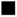 Spain; 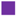 Ireland; 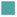 France; 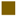 UK; and 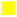 Austria.

Because the salt stress applied here was relatively mild, RGB colour–based vegetative indices showed almost no differences between salt-stressed plants and their corresponding controls. This was also observed in GLI and cive, the indices that appeared relevant biomarkers for low nutrition (Fig. 8E-F). However, the relative abundance of certain colour classes differed among genotypes: dark green (30:45:14) was reduced in varieties such as Maris Piper and Monalisa, whereas beige or light brown (154:147:126) accumulated in certain varieties such as Carlita (Fig. 8G-H).

Chlorophyll fluorescence-related parameters also did not differ significantly among varieties. PCA, representing PC1 and PC2 and accounting for 69.1% of the total variance, clustered the varieties similar to the distribution observed under low nutrition (Supplementary S5B). The biggest differences were observed in ETR, with Maris Piper and Nicola showing the greatest reduction in ETR4 (Fig. 8I, Supplementary S5B). Interestingly, Bermuda also tended to reduce ETR4 and presented the highest NPQ values under salt stress, parameters that were opposite correlated (Fig. 8I-J, Supplementary Fig. 5C). ETR4 was found to be positively correlated with both plant height and area (Supplementary Fig. S5B ), so its reduction could indicate that the electron transport obtained for light harvest was altered under salt stress, due to inhibition of chlorophyll biosynthesis, damage to the photosynthetic apparatus or restriction of the electron flow from PSII to PSI (Vineeth *et al*., 2023; Zahra *et al*., 2023).

Together, these results show that LOCOPOTS can detect genotype-dependent responses to moderate salt stress within three weeks using a combination of growth, colour and fluorescence traits. Under salinity, growth-related traits, particularly final plant height and Area_AUC, were more sensitive than broad colour indices or most fluorescence parameters. However, fluorescence traits such as ETR4 and NPQ provided complementary information on photosynthetic electron transport and photoprotective energy dissipation, helping to distinguish genotypes with similar growth responses but different physiological strategies.

### Salt stress reprogrammes carbon- and nitrogen-related metabolism in a genotype-dependent manner

To further understand the contrasting morphological and physiological responses observed under salinity stress, we selected seven genotypes that differed in their responses to 50 mM NaCl. The selected genotypes included Gala, which maintained growth under salt stress and was positioned in the tolerant/less-affected region of the PCA; Beltza, which showed a distinct metabolic profile and was clearly separated from the other genotypes; Nicola, which showed reduced growth and altered fluorescence performance under salt stress; BF-15, which showed a response similar to Nicola but with less pronounced changes in fluorescence-related parameters; Avano and Bermuda representing slow-growing genotypes under both control and salt stress conditions; and Bintje, which showed intermediate behaviour compared with the other selected genotypes (Figure 9). These genotypes displayed contrasting metabolic adjustments, indicating that potato responses to salt stress involved genotype-specific reprogramming of carbon and nitrogen metabolism.

**Fig. 9.**
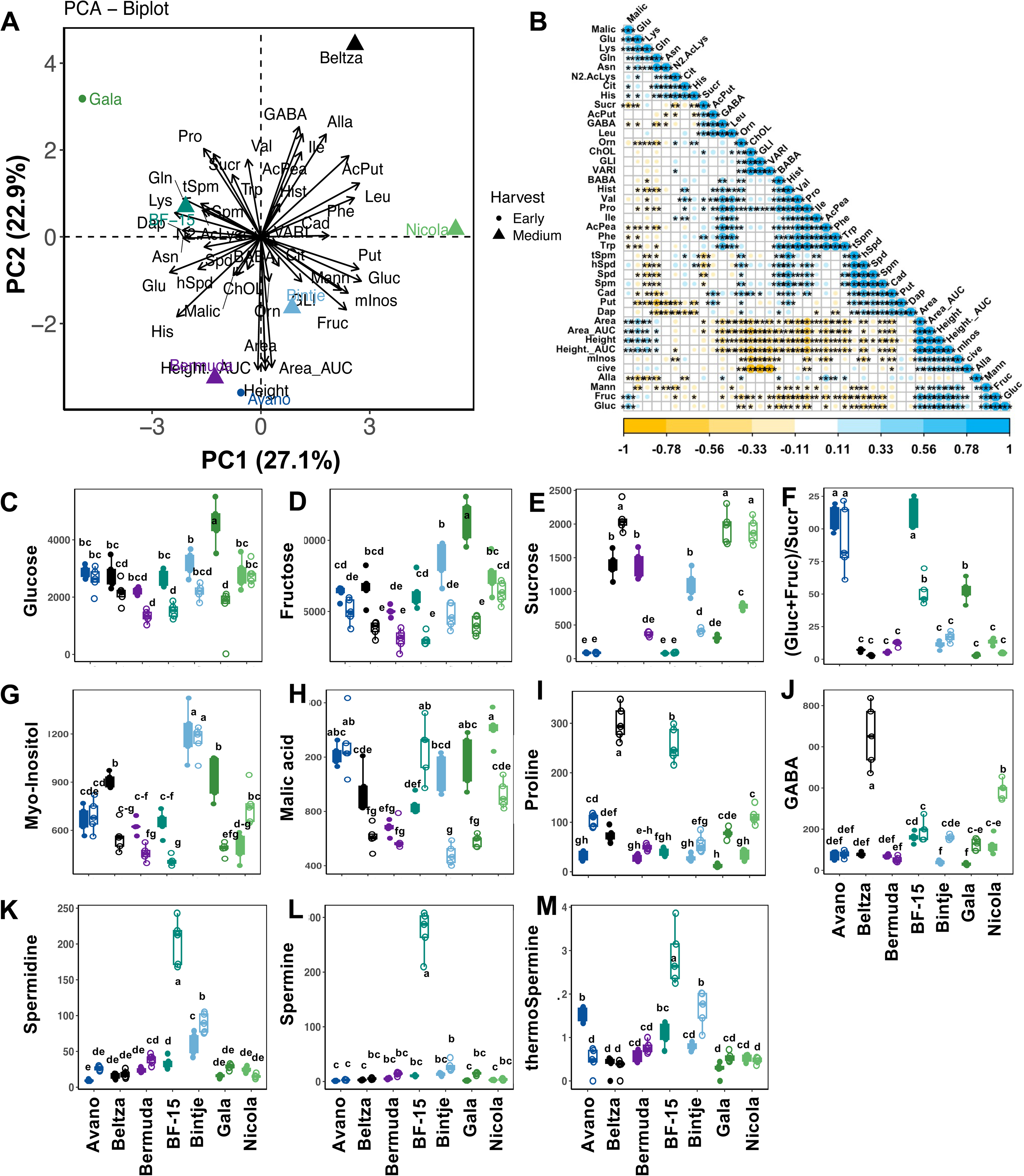
Biochemical changes in potato genotypes grown *in vitro* under salt stress conditions. **(A)** Principal component analysis (PCA) performed using sugars, free amino acids, free and acetylated polyamines, and other stress-related metabolites, together with representative RGB-derived traits. Values were expressed as the ratio between plants grown under salinity, using full-strength MS medium supplemented with 50 mM NaCl, and those grown on full-strength MS medium, 21 days after explant transfer. **(B)** Correlation matrix showing relationships between metabolite content and selected RGB-derived traits. Representative metabolites contributing to genotype separation are shown, including glucose (Gluc) **(C)**, fructose (Fruc) **(D)**, sucrose (Suc) **(E)**, the (Gluc + Fruc)/Suc ratio **(F)**, myo-inositol (mInos) **(G)**, malic acid (Malic) **(H)**, proline (Pro) **(I)**, γ-aminobutyric acid (GABA) **(J)**, spermidine **(K)**, spermine **(L)**, and thermospermine (tSpm) **(M)**. Different letters indicate significant differences according to Tukey’s HSD test following two-way ANOVA with genotype and treatment as factors. Filled boxplots and symbols represent plants grown on full-strength MS medium, whereas open boxplots and symbols represent plants grown under salt stress. Colours indicate genotype origin: 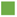 Germany; 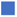 Netherlands; 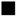 Spain; 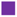 Ireland; 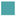 France. Additional metabolite abbreviations: 1,3-diaminopropane (Dap), allantoic acid (Alla), N^2^-acetyllysine (N2AcLys), acetylphenylethylamine (AcPea), acetylputrescine (AcPut), asparagine (Asn), β-aminobutyric acid (BABA), cadaverine (Cad), citrulline (Cit), glutamic acid (Glu), glutamine (Gln), histamine (Hist), histidine (His), homospermidine (hSpd), isoleucine (Ile), leucine (Leu), lysine (Lys), mannitol (Mann), ornithine (Orn), phenylalanine (Phe), proline (Pro), putrescine (Put), tryptophan (Trp), and valine (Val).

PCA analysis, in which PC1 and PC2 explained 50.0% of the total variance, separated the genotypes according to their metabolic and phenotypic responses to salinity (Fig. 9A). Gala was in the upper-left part of the biplot, with BF-15 nearby, and was correlated to Pro, Gln, Spm and tSpm. In contrast, Bintje, located opposite to them, was associated with higher content of monosaccharides such as glucose and fructose, as well as the polyols mannitol and myo-inositol. On the negative side of PC1, which was associated with higher growth-related traits, including plant height and Area_AUC, were Bermuda and Avano, whereas Nicola was positioned close to the x-axis of PC2. Beltza was located opposite Bermuda and Avano and was associated with higher GABA levels, among other metabolites (Fig. 9A). This distribution suggests that the maintenance of growth under salt stress was not linked to a single metabolic strategy, but rather to different genotype-dependent adjustments in primary and secondary metabolism.

Among the measured carbohydrates, glucose and fructose showed contrasting genotype-dependent patterns under salt stress (Fig. 9C-D). In Gala, maintaining growth under salinity was associated with marked reductions in glucose and fructose, whereas in Nicola, these monosaccharides did not change. Interestingly, Gala and Bintje exhibited the highest levels of glucose and fructose under control conditions, but decreased under salinity (Fig. 9C-D). Soluble sugars are commonly involved in osmotic adjustment under salt stress and can contribute to cellular protection by stabilising membranes and proteins, maintaining osmotic balance and supporting energy metabolism (Munns and Tester, 2008; Krasensky and Jonak, 2012). In this regard, the association between growth-related traits and monosaccharide levels suggests that the ability to regulate soluble sugar availability under salt stress may help sustain growth and carbon metabolism *in vitro*. However, the high monosaccharide content observed under optimal conditions in Bintje and Gala, together with the reduction of these sugars to levels similar to those of other genotypes under stress, and the high myo-inositol content in both genotypes, suggests that full-strength MS medium containing 3% sucrose may not be equally optimal for all potato genotypes. Moreover, differences in sucrose and in the (glucose + fructose)/sucrose ratio indicate that each genotype responded to salt stress through distinct adjustments in carbon metabolism (Fig. 9F). Salt stress is known to alter sucrose metabolism by shifting the balance among sucrose synthesis, cleavage, and transport, thereby affecting source–sink relationships and growth. In this context, the ratio between glucose plus fructose and sucrose may provide additional information on carbohydrate remobilisation (Fig. 9F). Higher values of this ratio may reflect increased cleavage of sucrose into hexoses, providing readily available carbon skeletons and osmotic adjustment capacity under salinity. Interestingly, Avano and BF-15 maintained the highest values of this ratio, but both were low-growing genotypes, suggesting that a high (glucose + fructose)/sucrose ratio may also reflect growth limitation or altered carbon use rather than improved salt tolerance.

Polyols and compatible solutes also differed among genotypes. Myo-inositol and malic acid showed genotype-specific patterns under salt stress (Fig. 9G-H). Bintje showed the highest myo-inositol content under both control and salt stress conditions, whereas Avano, BF-15 and Nicola had the highest levels under salinity. As discussed above, myo-inositol is involved in osmotic adjustment, redox balance, and stress protection, and it also acts as a precursor for phosphoinositides and other signalling molecules (Valluru and Van den Ende, 2011). However, its accumulation was not correlated with the biomass-related parameters, suggesting that the presence of these metabolites may indicate stress perception or acclimation rather than tolerance per se.

Amino acid metabolism was also strongly affected by salinity. Pro accumulated significantly, particularly in Beltza, BF-15 and Nicola (Fig. 9I). Pro is one of the most widely reported compatible solutes under salt and osmotic stress, contributing to osmotic adjustment, ROS detoxification, and protection of cellular structures (Szabados and Savouré, 2010). However, Pro accumulation is often greater in plants under higher stress and does not necessarily indicate improved growth performance (De Diego *et al*., 2012). In our dataset, the association of proline with genotypes showing stronger growth reduction suggests that Pro may reflect stress intensity rather than salt tolerance under these conditions.

GABA also contributed to the separation of genotypes, with higher values in Beltza and Nicola (Fig. 9J). GABA metabolism is closely linked to carbon–nitrogen balance, pH regulation, stress signalling and the tricarboxylic acid cycle through the GABA shunt (Michaeli and Fromm, 2015; Podlešáková *et al*., 2019). Its accumulation under salt stress may, therefore, indicate metabolic reconfiguration to maintain energy production and nitrogen balance when primary metabolism is constrained. The strong association of GABA with Beltza and Nicola, which were among the most affected genotypes in fluorescence-related parameters (Fig. 8I-J), suggests that these genotypes activated stress-related nitrogen metabolism and alternative carbon metabolism more strongly than Gala did. Under conditions in which carbon availability or carbon use is limited, plants can redirect carbon from amino acid pools back into respiration via the GABA shunt to sustain primary metabolism (Ji *et al*., 2020; Saiz-Fernández *et al*., 2020).

Salt stress also modified polyamine metabolism (Fig. 9K–M). Under salinity, Spd, Spm and tSpm were mainly accumulated in BF-15. Polyamines are involved in membrane stabilisation, ROS homeostasis, ion balance, and stress signalling, and their accumulation has often been associated with improved tolerance to salinity and other abiotic stresses (Alcázar *et al*., 2010; Tiburcio *et al*., 2014). However, in our data, high polyamine accumulation was not necessarily associated with the best growth performance; in fact, the opposite was observed. This suggests that, under the moderate salt stress imposed here, polyamine accumulation may reflect the activation of protective responses in more severely affected genotypes (sensitive) rather than serving as a direct marker of maintained growth, or indicative of plant tolerance.

Overall, these results indicate that salt stress induced a strong genotype-dependent reprogramming of carbon- and nitrogen-related metabolism in *in vitro* potato plants. Genotypes that maintained growth, such as Gala, did not necessarily accumulate the highest levels of classical stress metabolites. In contrast, genotypes with stronger growth or fluorescence impairment, such as Beltza, BF-15, and Nicola, showed greater accumulation of Pro, GABA, polyamines, or allantoic acid, suggesting activation of stress-response and nitrogen-remobilisation pathways. Thus, under 50 mM NaCl, the accumulation of stress-related metabolites appeared to reflect stress intensity and metabolic acclimation more than salt tolerance per se. These findings highlight the value of combining LOCOPOTS-derived phenotypic traits with targeted metabolite profiling to distinguish genotypes that maintain growth under salinity from those that respond by activating stronger protective metabolism.

## Conclusion

This study presents LOCOPOTS, a low-cost, compact and modular pipeline for high-throughput phenotyping of *in vitro* potato plants under control and abiotic stress conditions. By combining optimised *in vitro* culture in individual square vessels, standardised RGB image acquisition, automated U-Net-based segmentation, and trait extraction, LOCOPOTS overcomes several limitations associated with conventional potato phenotyping, particularly the high space requirements, costs, and handling time of potted-plant platforms. The system was reproducible across four independent experimental rounds performed over more than one year, supporting its robustness for repeated phenotyping campaigns.

The comparison among manual, semi-automatic and automatic image-analysis approaches demonstrated that our in-house Python pipeline reliably extracted plant height and growth-related traits while reducing post-processing time and avoiding segmentation artefacts observed with semi-automatic analysis. Growth-integrated traits, particularly Area_AUC, was more informative than single time-point measurements because they captured the dynamics of plant development throughout the 21-day phenotyping period. In addition, RGB-derived colour classes and vegetation indices provided complementary information on plant greenness, pigmentation, and potential physiological status, allowing rapid discrimination among potato genotypes grown under control conditions.

LOCOPOTS also detected genotype-dependent responses to low nutrient availability and salt stress. Under low-nutrient conditions, Bintje showed improved growth and a distinctive physiological and metabolic profile, including changes in GLI, cive, fructose, BABA, Spd, N^2^-acetyllysine, myo-inositol, and allantoic acid. These results suggest that low nutrient responses were not driven by a single trait, but by coordinated reprogramming of carbon and nitrogen metabolism. Importantly, the reduction of myo-inositol in genotypes such as Bintje and Gala highlights that full-strength MS medium supplemented with 3% sucrose may not be equally optimal for all potato genotypes, and that culture medium composition can strongly influence in vitro growth and metabolic status.

Under moderate salt stress, growth-related RGB traits, particularly final plant height and Area_AUC, were more sensitive than broad colour indices or most chlorophyll fluorescence parameters. Genotypes that maintained growth under salinity did not necessarily accumulate the highest levels of classical stress metabolites. In contrast, genotypes with stronger growth or fluorescence impairment showed greater accumulation of Pro, GABA and polyamines, suggesting that these metabolites mainly reflected stress intensity and metabolic acclimation rather than tolerance per se. Thus, integrating LOCOPOTS-derived phenotypic traits with targeted metabolite profiling enabled the distinction between genotypes that maintained growth under stress and those that activated stronger protective or stress-response metabolism.

Overall, LOCOPOTS provides a scalable platform for early-stage screening of potato genetic diversity under defined *in vitro* stress conditions. Its low cost, small spatial footprint and compatibility with automated image analysis make it particularly suitable for breeding programmes, pre-breeding studies and genotype maintenance in potato genebanks. Although *in vitro* phenotyping cannot replace field or greenhouse validation, it offers a powerful first-screening layer for identifying contrasting genotypes, prioritising candidates for further testing, and generating hypotheses about the physiological and metabolic mechanisms underlying abiotic stress responses in potato.

Future applications of LOCOPOTS could include large-scale screening of diverse germplasm collections, testing of stress-alleviating compounds or biostimulant products, and identification of genotype × treatment combinations that improve performance under defined stress conditions. In addition, this platform could support the optimisation of *in vitro* culture protocols for the storage, maintenance and propagation of recalcitrant or problematic genotypes, helping to establish more genotype-tailored standards for potato genebanks and breeding pipelines.

## Supplementary data

The following supplementary data are available online at *JXB*.

**Fig. S1.** Potato varieties used in this study and their origin.

**Fig. S2.** Low-cost in-house system for RGB imaging of potato plants grown *in vitro*.

Fig. S3. RGB Image-derived morphological and colour traits, and ChlF-derived traits of potato genotypes grown *in vitro* under control conditions. (A) Correlation matrix performed using image-derived traits from 30 potato genotypes grown on full-strength MS medium for 21 days after explant transfer. The analysed features included four morphological traits — plant height, projected area, height AUC and area AUC — together with 15 segmentation-derived colour parameters and 50 colour-based vegetation indices. **(B)** Principal component (PC) analysis using the morphological traits and the chlorophyll fluorescence-related parameters. **(C)** Correlation matrix performed using morphological and ChlF imaging-derived traits. 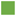 Germany; 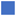 Netherlands; 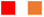 the USA; 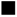 Spain; 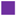 Ireland; 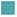 France; 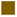 UK; and 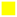 Austria.

Fig. S4. RGB Image-derived morphological and ChlF-derived traits of potato genotypes grown *in vitro* under low nutrient conditions. (A) Representative genotype-specific distributions are shown for selected RGB-derived traits contributing to genotype separation, including final plant height. (B) Correlation matrix performed using ChlF image-derived traits from 30 potato genotypes grown on low nutrient conditions for 21 days after explant transfer. 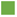 Germany; 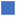 Netherlands; 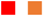 the USA; 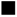 Spain; 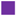 Ireland; 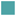 France; 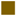 UK; and 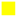 Austria.

Fig. S5. RGB Image-derived morphological and ChlF-derived traits of potato genotypes grown *in vitro* under salt stress conditions. (A) Correlation matrix performed using image-derived traits from 30 potato genotypes grown on full-strength MS medium for 21 days after explant transfer. The analysed features included four morphological traits — plant height, projected area, height AUC and area AUC — together with 15 segmentation-derived colour parameters and 50 colour-based vegetation indices. **(B)** Principal component (PC) analysis using the morphological traits and the chlorophyll fluorescence-related parameters. **(C)** Correlation matrix performed using ChlF image-derived traits from 30 potato genotypes grown on low nutrient conditions for 21 days after explant transfer. 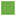 Germany; 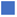 Netherlands; 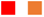 the USA; 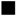 Spain; 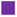 Ireland; 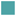 France; 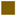 UK; and 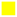 Austria.

**Table S1.** Equations used for calculating the RGB-based vegetation indexes.

**File S1.** ANOVAs and post-hoc analysis using Tukey’s HSD test.

## Author contributions

I.S.F., L.S. and N.D.D. led the idea and design of the experimental work. I.S.F., L.S. and N.D.D. conceived the manuscript’s structure. J.I.R.D.G., M.L., and A.O.B. provided the original plants and contributed to establishing the *in vitro* culture method. I.S.F. and L.A.B.P. contributed to the plant phenotyping experiments and image acquisition. P.K. contributed to the experimental work regarding the machine learning model and the overall image analysis pipeline. I.S.F. and L.A.B.P. performed the manual analysis of the phenotyping imaging. S.Č.Z performed the metabolomic analysis. I.S.F., L.A.B.P. and N.D.D performed the data analysis. I.S.F., L.A.B.P. and N.D.D. contributed to the writing of the manuscript. All authors reviewed and agreed on the manuscript before submission.

## Data availability

All data used in this study, including the codes and a readme for their use, are openly available on ZENODO (10.5281/zenodo.20122647).

---

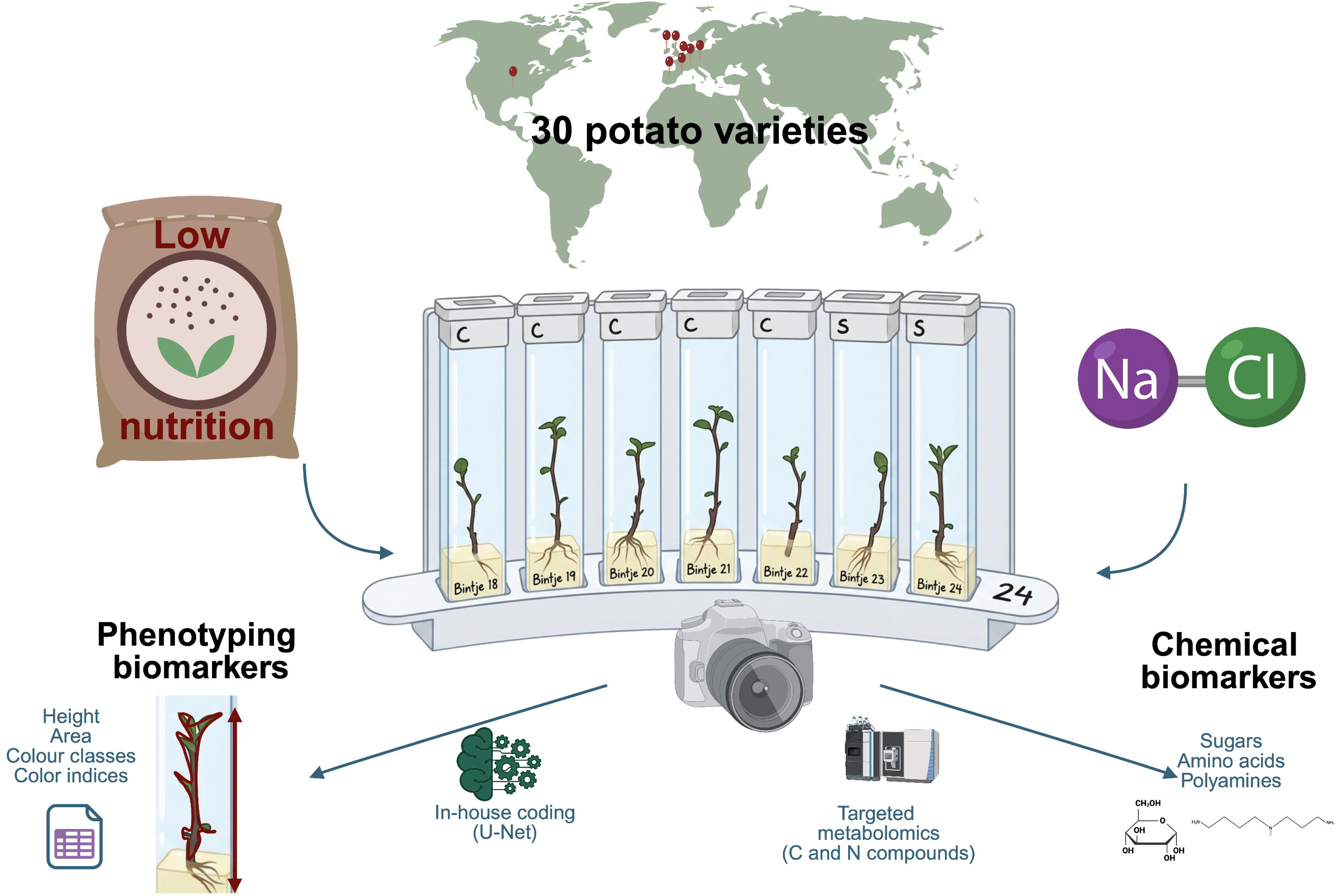

